# High Diversity Gene Libraries Facilitate Machine Learning Guided Exploration of Fluorescent Protein Sequence Space

**DOI:** 10.64898/2026.03.01.706892

**Authors:** Anissa Benabbas, Phillip Kearns, Avery Billo, Lauren O. Chisholm, Calin Plesa

## Abstract

While protein language models (PLMs) have shown great promise for protein design, their performance is fundamentally constrained by the diversity and completeness of available training data. In particular, PLMs often struggle to extrapolate to sequences that fall outside the distribution spanned by their training sets, limiting their ability to discover proteins in sparsely sampled regions of sequence space. Here we test the hypothesis that experimentally expanding training diversity can convert extrapolation into interpolation and thereby enable discovery of functional sequences beyond natural protein manifolds. Using large-scale gene synthesis and DNA shuffling, we generate libraries that span a broad region of fluorescent protein sequence space and create chimeric variants that bridge between distant homologs. Functional screening for blue fluorescence yields thousands of active variants distributed across diverse sequence lineages. Fine-tuning ProtGPT2 on this expanded dataset enables generation of diverse fluorescent proteins, including designs that extend beyond the regions occupied by known natural sequences while retaining function. This work illustrates how synthetic approaches can help address key limitations in machine learning-guided protein design, especially for small or sparsely populated protein families, by actively creating novel sequences across unexplored but functional regions of sequence space.

## Introduction

The advent of machine learning (ML) technologies has introduced a paradigm shift in how we approach protein engineering. Recent breakthroughs have demonstrated remarkable success in identifying patterns within sequence data and predicting protein structures^1,2^. These methods can prioritize limited experimental resources toward the most promising sequences and can even generate entirely novel sequences distant from any natural homolog^3–7^. The appeal of ML lies in its ability to infer key sequence-function relationships without detailed knowledge of the underlying biophysical mechanisms. Unlike traditional directed evolution, ML-guided approaches do not require a smooth local fitness landscape and can, in principle, access regions of sequence space that are inaccessible by stepwise evolutionary trajectories^8^. Because the primary costs of ML are computational, they are often modest relative to the experimental burden of large-scale screening. Generative models have now been used to make novel sequences unseen in nature^7,9–16^, including sequences that are highly divergent from their training data^17,18^, some of which are functional^6,19–22^. Parallel efforts aim to couple generative models with predictive models to allow for optimization (design) of specific protein properties^23–28^.

Despite this progress, a central bottleneck remains in the composition of the training data itself. ML models perform best when predicting within the distribution of sequences they have observed. When asked to generalize beyond that distribution, performance often declines substantially. Thus, the diversity and structure of the training set can be as important as the model architecture. Recent work consistently shows that ML models struggle with extrapolation relative to interpolation^29–32^. Extrapolation refers to predictions made on data that fall outside the distribution spanned by the training set, whereas interpolation refers to predictions within that distribution (Fig. 1A-B). In the context of protein sequence space, this distinction is critical. If the training set sparsely samples a protein family, many candidate sequences lie outside the sampled manifold and require extrapolation. By contrast, if the training data densely and broadly sample sequence space, many previously unseen sequences become interpolative queries, reducing model uncertainty and increasing predictive reliability (Fig. 1A). This distinction has important implications for protein engineering. The highest-fitness genotype for a desired phenotype is often not a variant of the most studied homolog. Instead, global fitness optima may reside in distant regions of sequence space that are poorly connected to well-studied laboratory templates. When engineering efforts narrowly focus on mutagenizing a single parent protein, they are effectively confined to local peaks in the fitness landscape. Expanding the diversity of starting genotypes increases the probability of identifying higher fitness maxima that would otherwise remain inaccessible. However, generating sufficiently large and diverse datasets remains a major challenge, both for ML model training^21,33^ and for initiating directed evolution campaigns. The relationship between training set size and ML model performance is well-documented, with larger datasets generally yielding improved classification and regression accuracy^34,35^. The required data volume also depends on the complexity of the underlying sequence–function relationships^36^. In proteins, this complexity arises from multiple effects, including epistatic interactions between residues that may be distant in primary sequence yet coupled in structure and function. Importantly, increasing sample size alone is insufficient if the added sequences are highly redundant or confined to a narrow mutational neighborhood. True improvement requires expansion of both scale and diversity.

**Fig. 1.**
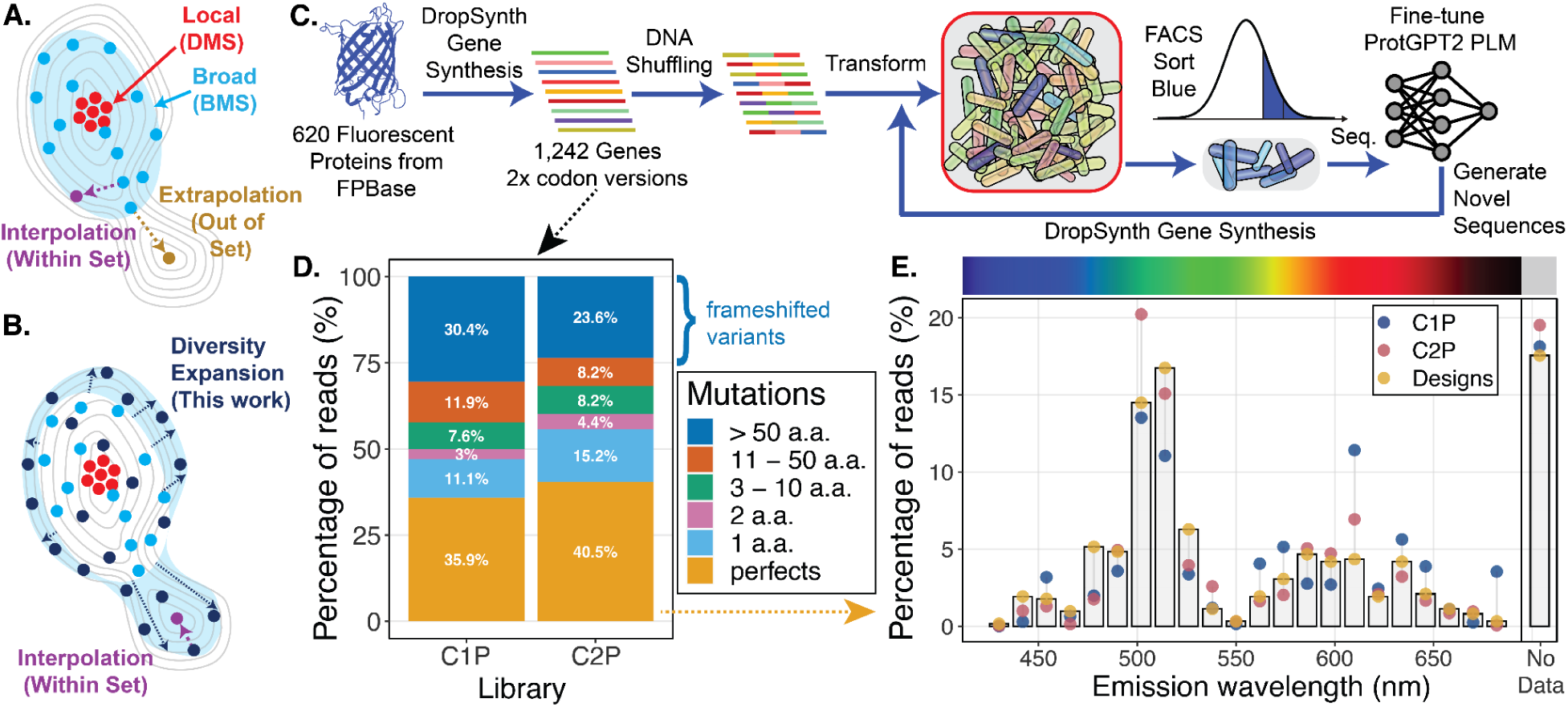
DropSynth Assembly of Fluorescent Protein Libraries. **A.** Conceptual illustration of interpolation versus extrapolation in protein sequence space. Deep mutational scanning (DMS) probes local neighborhoods around a parent sequence, while broad mutational scanning (BMS) of related homologs extends further out. For protein language models trained on such data, predictions within the sampled region correspond to interpolation and are generally more reliable than extrapolative predictions outside the training distribution. **B.** Diversity expansion strategy used in this study. By synthesizing and recombining a wide set of homologous fluorescent proteins, we intentionally expand the sampled sequence manifold. This approach aims to convert many previously extrapolative prediction problems into interpolative ones by increasing the density and breadth of experimentally characterized sequences. **C.** Experimental workflow. A set of 620 fluorescent proteins was synthesized in two synonymous codon versions using DropSynth. The resulting parental libraries were subjected to DNA shuffling to generate chimeric diversity, transformed, and screened by fluorescence-activated cell sorting for blue emission. Selected variants were sequenced and used to fine-tune the ProtGPT2 protein language model. Model-generated sequences were subsequently synthesized and experimentally characterized, further expanding the functional training dataset. **D.** Distribution of assembly outcomes in parental libraries. At least 35% of reads in each library correspond to perfect amino acid sequences matching the intended design. More than 20% of reads represent low-distance mutants containing 1 to 10 amino acid substitutions. A large fraction of the remaining reads consist of higher-distance frameshifted variants. **E.** Spectral representation of assembled libraries. The percentage of reads corresponding to perfect sequences at each emission wavelength is shown for C1P (blue) and C2P (red) relative to the designed distribution (gold points and gray bars). The observed enrichment of green-emitting proteins mirrors the underlying FPBase distribution.

To effectively model relationships between amino acid sequence and quantitative properties such as fluorescence brightness, training datasets must span a broad region of functional sequence space. Traditional mutagenesis methods including error-prone PCR, nicking mutagenesis, and deep mutational scanning have been powerful tools for protein optimization. However, they primarily generate variants that retain close homology to a parent template, limiting the accessible diversity. Such approaches enable detailed mapping of local fitness landscapes, but rarely expand the global boundaries of a protein family. As a result, many ML-guided design tasks remain fundamentally extrapolative. Although data augmentation^37^ has expanded effective dataset size in fields such as computer vision and natural language processing^38^, its application to biological sequences^39^ is constrained by functional requirements. Unlike images or text, protein sequences cannot be arbitrarily perturbed without disrupting folding or activity. Functional diversity must therefore be validated experimentally rather than synthetically through trivial transformations.

Overcoming these limitations requires functionally characterized datasets that combine scale with broad sequence dispersion. While advances in sequencing have produced vast quantities of metagenomic data, most sequences remain uncharacterized and unevenly distributed across protein families. Recent planetary-scale assembly efforts such as LOGAN^40^ have demonstrated that public sequencing archives contain orders of magnitude more protein diversity than currently represented in curated databases, yet the vast majority of these sequences lack functional characterization. Furthermore, many protein families of interest exhibit limited natural diversity. Even for well-studied systems, deep mutational scanning only explores local neighborhoods around selected parental sequences. In contrast, large-scale gene synthesis technologies like DropSynth^41–43^ offer access to both natural diversity and designed variants. By assembling thousands of full-length genes in parallel, these methods allow deliberate expansion of sequence space while preserving functional constraints. Synthetic recombination strategies, including DNA shuffling, can further increase sequence dispersion by combining diverse homologs into novel chimeras.

We hypothesized that generating highly diverse, functionally characterized training data would convert many extrapolative ML prediction problems into interpolative ones. By expanding the sampled sequence manifold of a protein family, previously distant regions of sequence space can be brought within the effective scope of the model (Fig. 1A-B). To test this hypothesis, we focused on β-barrel fluorescent proteins as a model system. Fluorescent proteins provide a direct and quantitative functional readout, making them an ideal platform for integrating gene synthesis, high-throughput screening, and ML-guided design. Since the discovery of Aequorea victoria green fluorescent protein^44,45^ (avGFP), systematic searches for natural homologs^46–50^ and significant protein engineering efforts^51,52^ have expanded the family. Nonetheless, the total number of β-barrel fluorescent proteins remains limited, with fewer than 800 entries cataloged in databases such as FPBase^53^. This relatively small family size constrains the natural sequence diversity available for ML training. Although several deep mutational scans^54–56^ have mapped local fitness landscapes of specific fluorescent proteins, these studies provide only partial coverage of global sequence space. Ongoing engineering efforts seek brighter variants^57–59^, particularly in the blue and red^60,61^ spectral regions, as well as improved photostability^62^ for imaging applications, or build fluorescent biosensors^63^. Previously, ML approaches have been applied to fluorescent protein optimization^64^ using diverse strategies^65^, including 3D convolutional neural networks^66^, biophysics-based models^67^, small libraries of random mutants^21^, and guided miniaturization^68^. However, these studies mostly operate within limited regions of sequence space defined by a small number of parental templates.

In this study, we synthesized gene libraries encompassing nearly all known β-barrel fluorescent proteins, and applied DNA shuffling to generate highly diverse chimeric libraries (Fig. 1C). This strategy intentionally broadened the functional sequence manifold of the family, enabling exploration of combinations not sampled by evolution or single-template mutagenesis. Using fluorescence-activated cell sorting and next-generation sequencing, we identified functional blue fluorescent proteins from these libraries and assembled a large, diverse training dataset. We then fine-tuned a generative ML model, ProtGPT2, on this expanded dataset and used it to design 1,536 novel blue fluorescent proteins, which were synthesized and experimentally characterized. By coupling high-diversity gene synthesis with iterative ML-guided design, we aim to show that deliberate expansion of sequence diversity reduces reliance on extrapolation and increases the probability of identifying higher-fitness optima within a small protein family.

## Results and Discussion

### DropSynth Assembly and Characterization of Parental Fluorescent Protein Libraries

We first sought to construct a broadly representative map of fluorescent protein sequence space. To do this, we collected 620 β-barrel fluorescent protein sequences from FPBase and synthesized them using DropSynth in two independent codon-optimized versions, yielding 1,242 unique gene constructs. Each amino acid sequence was represented with two synonymous codon variants, generating two parental libraries, C1P and C2P (Fig. S1). To enable pooled functional analysis, these libraries were cloned into a high-copy, barcoded expression vector derived from pUC19. Long-read sequencing revealed coverage rates of 87.7% (544 of 620) and 79.8% (495 of 620) for the C1P and C2P libraries, respectively (Fig. S2A). When combined across both codon versions, total protein coverage increased to 94.0% (583 of 620), indicating that most parental amino acid sequences were successfully assembled in at least one codon context. The dual codon strategy increased effective amino acid coverage and mitigated synthesis biases associated with individual codon designs. Alignment of mapped reads to their nearest parental designs revealed a median percentage of perfect amino acid sequences of 35.7% (s.d. 12.7%) for C1P and 33.1% (s.d. 10.4%) for C2P (Fig. S2B) for proteins with at least 100 aligned barcodes. On a per-read basis, 35.9% and 40.5% of total reads corresponded to perfect amino acid sequences in C1P and C2P, respectively (Fig. 1D). An additional 21.7% and 27.8% of reads were low-distance mutants containing between 1 and 10 amino acid substitutions. The remaining reads consisted primarily of higher distance frameshifted sequences. Notably, the substantial fraction of low-distance mutants provides additional local diversity that may be informative for ML training, while the perfect sequences preserve the intended global diversity of the FPBase set. Library uniformity was evaluated using the Gini coefficient, yielding values of 0.68 for C1P and 0.64 for C2P, comparable to previous DropSynth-assembled libraries. Although some representation bias remains, the observed distribution supports sufficiently broad sampling for high-throughput functional screening and ML training. We next examined whether assembled libraries preserved the intended distribution of spectral diversity. The percentage of reads corresponding to perfect sequences across emission wavelengths ranging from 424 nm to 698 nm is shown in Fig. 1E. The observed distribution closely tracks the designed distribution (gold color points and bars), with green-emitting proteins overrepresented relative to other colors, reflecting the underlying distribution of fluorescent proteins in FPBase rather than assembly bias.

### DNA Shuffling Creates Functional Chimeric Proteins

To further expand the diversity of our fluorescent protein libraries beyond the parental set, we applied DNA shuffling^69^ to generate large numbers of chimeric variants. DNA shuffling is a synthetic recombination strategy in which pooled genes are randomly fragmented using DNase I and subsequently reassembled by low-stringency PCR, enabling crossover between homologous sequences. This process creates mosaic genes that combine segments from multiple parental templates and can traverse regions of sequence space that are not accessible through point mutagenesis alone. We combined both parental libraries (C1P and C2P) prior to shuffling to maximize starting-template diversity, generating the shuffled library C12S. Because the parental set already spans hundreds of homologous β-barrel fluorescent proteins, recombination between these templates provides a combinatorial expansion of sequence diversity that substantially exceeds the original design space. PacBio sequencing of the shuffled library revealed a large increase in diversity. When normalized for sequencing depth, the number of unique protein variants observed in the shuffled library was approximately threefold higher than the combined parental libraries (Fig. 2A). This increase reflects the combinatorial generation of novel sequence combinations rather than incremental mutational drift. Only 2.2% of unique protein variants detected in the parental libraries were also present in the shuffled library (Fig. 2B), demonstrating that recombination largely displaced the original sequence population. At a raw read depth of 728,479 reads, we identified 87,436 unique shuffled protein sequences that were not observed in either parental library (Fig. 2B). These results confirm that DNA shuffling substantially expands the functional sequence manifold rather than merely reshuffling representation frequencies.

**Fig. 2.**
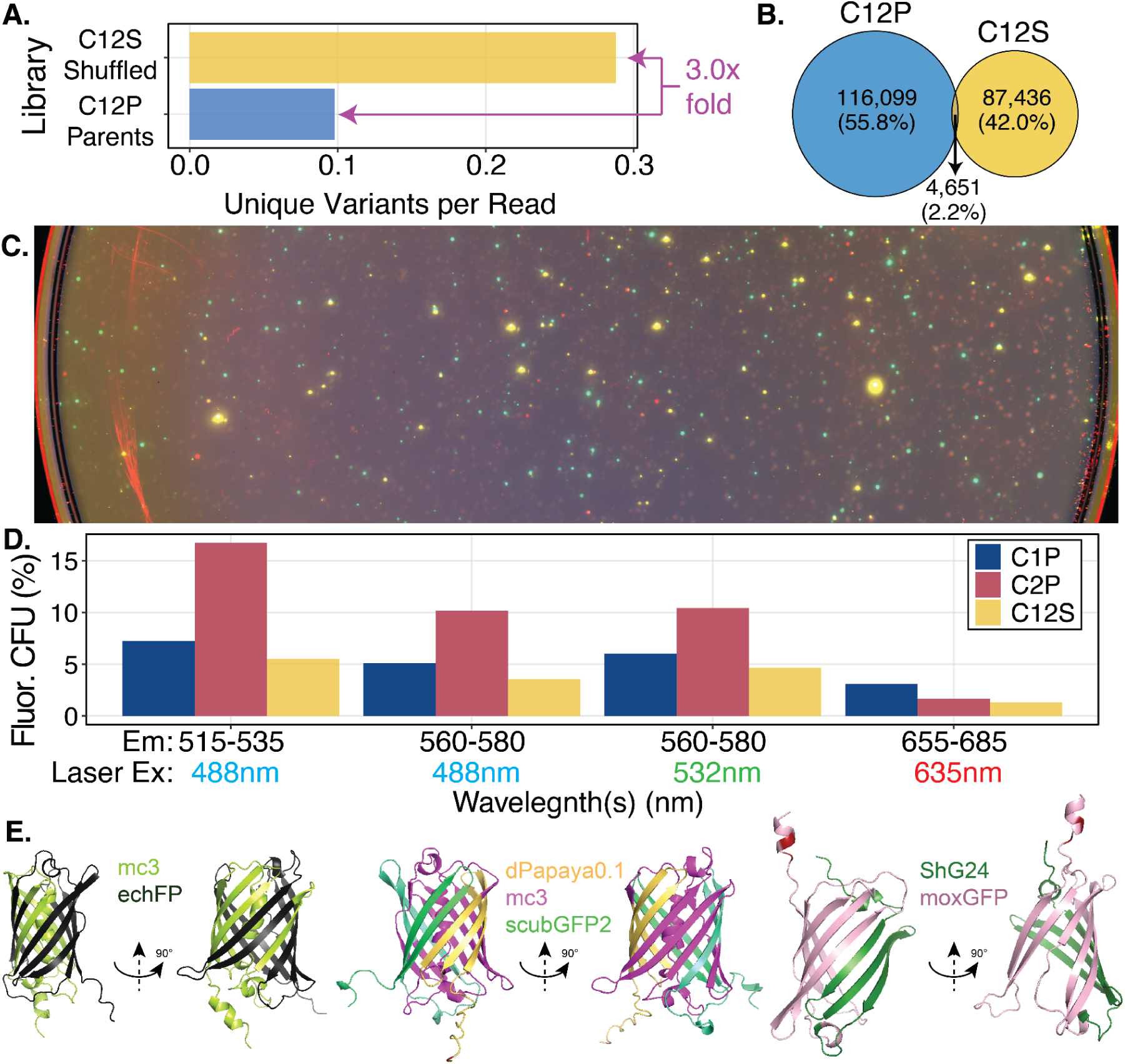
DNA Shuffling Expands Sequence Diversity While Retaining Functional Fluorescence. **A.** Diversity expansion following DNA shuffling. The number of unique protein variants normalized by sequencing depth shows a threefold increase in the shuffled library (C12S) relative to the combined parental libraries (C12P). Values are normalized as unique amino acid sequences per read to account for differences in sequencing depth. **B.** Overlap between parental and shuffled libraries. Only 2.2% of unique parental variants are also observed in the shuffled library. Over 87,400 unique protein variants were detected exclusively in C12S at a raw read depth of 728,479, demonstrating that recombination generates largely novel sequence combinations. **C.** Functional fluorescence of shuffled variants. Representative false-color overlay of Typhoon laser scanner images of plated colonies expressing shuffled fluorescent proteins under multiple excitation and emission conditions. Fluorescent colonies are observed across a range of spectral channels, indicating retention of functional chromophore formation following recombination. **D.** Fraction of fluorescent colony forming units (CFUs) across spectral channels. The percentage of fluorescent CFUs is shown for parental libraries (C1P and C2P) and the shuffled library (C12S) under different excitation and emission filter combinations. Although shuffling reduces the overall fraction of fluorescent colonies relative to parental libraries, substantial functional diversity remains, reflecting the structural tolerance of the β-barrel scaffold to segmental recombination. **E.** Structural mapping of representative chimeric proteins. Amino acid alignments of shuffled variants to parental templates are mapped onto predicted structures, with parental fragment origins color-coded. Distinct mosaic architectures illustrate recombination of structural segments across the β-barrel fold, confirming successful generation of chimeric proteins.

We next assessed whether this increase in diversity was accompanied by retention of functional fluorescence. Colonies expressing the parental libraries (C1P and C2P) and the shuffled library (C12S) were plated and imaged using a Typhoon laser scanner with four excitation and emission combinations spanning different spectral regions. False-color overlays of representative scans are shown in Fig. 2C, with the uncropped image provided in Fig. S3. Fluorescent colony forming units were quantified from plate images to determine the fraction of functional variants in each library (Fig. 2D). Across the four excitation and emission settings, C1P exhibited a median of 5.6% fluorescent colony forming units (CFUs), while C2P exhibited 10.3%, consistent with the higher fraction of perfect sequences observed in C2P. The shuffled library C12S exhibited a median of 4.1% fluorescent colonies. Hence, while DNA shuffling reduced the fraction of functional variants by approximately 1.9-fold relative to the mean of both parental libraries, it simultaneously increased overall sequence diversity by at least threefold. Overall, the shuffled library retained a substantial number of functional fluorescent proteins despite extensive recombination, indicating that the β-barrel scaffold can tolerate considerable segmental exchange.

To confirm the chimeric nature of shuffled sequences, we aligned representative variants to their nearest parental templates and mapped parental fragment contributions onto AlphaFold2 predicted protein structures (Fig. 2E). These alignments reveal clear mosaic architectures, with discrete structural segments derived from different parental proteins. Additionally, the overall protein length distribution of shuffled variants closely matched that of the parental libraries. The shuffled library exhibited a median protein length of 218 amino acids (s.d. 78), compared to a median of 211 amino acids (s.d. 74) for the combined parental set (Fig. S4). This indicates that recombination preserved global structural constraints of the β-barrel fold, rather than generating truncated or aberrantly extended sequences. Taken together, these results demonstrate that DNA shuffling produces a large population of novel, structurally plausible, and partially functional chimeric fluorescent proteins.

### Blue Fluorescence FACS sort to Generate Training Data

Having expanded sequence diversity through DNA shuffling, we next sought to isolate variants exhibiting blue fluorescence for the generation of a functionally enriched training dataset. The shuffled library C12S was subjected to fluorescence-activated cell sorting (FACS) using excitation and emission settings optimized for blue fluorescent proteins. Two bins were defined at the upper end of the brightness distribution, as shown in Fig. 3A and Fig. S5. The brightest bin, designated BS4, represented the top 0.13% of the population, while the adjacent bin BS3 represented 1.3%. PacBio sequencing of the sorted populations revealed strong divergence from the unsorted shuffled library. Only 1% of sequences overlapped between BS4 and C12S, and 1.1% overlapped between BS3 and C12S. In contrast, the overlap between BS3 and BS4 was 8.5%, as shown in Fig. 3A. This indicates that FACS strongly reshaped the sequence distribution, enriching a small subset of variants from a highly diverse background. The relatively modest overlap between BS3 and BS4 further suggests that even within the enriched fraction, brightness-based selection partitions sequence space into partially distinct subpopulations.

**Fig. 3.**
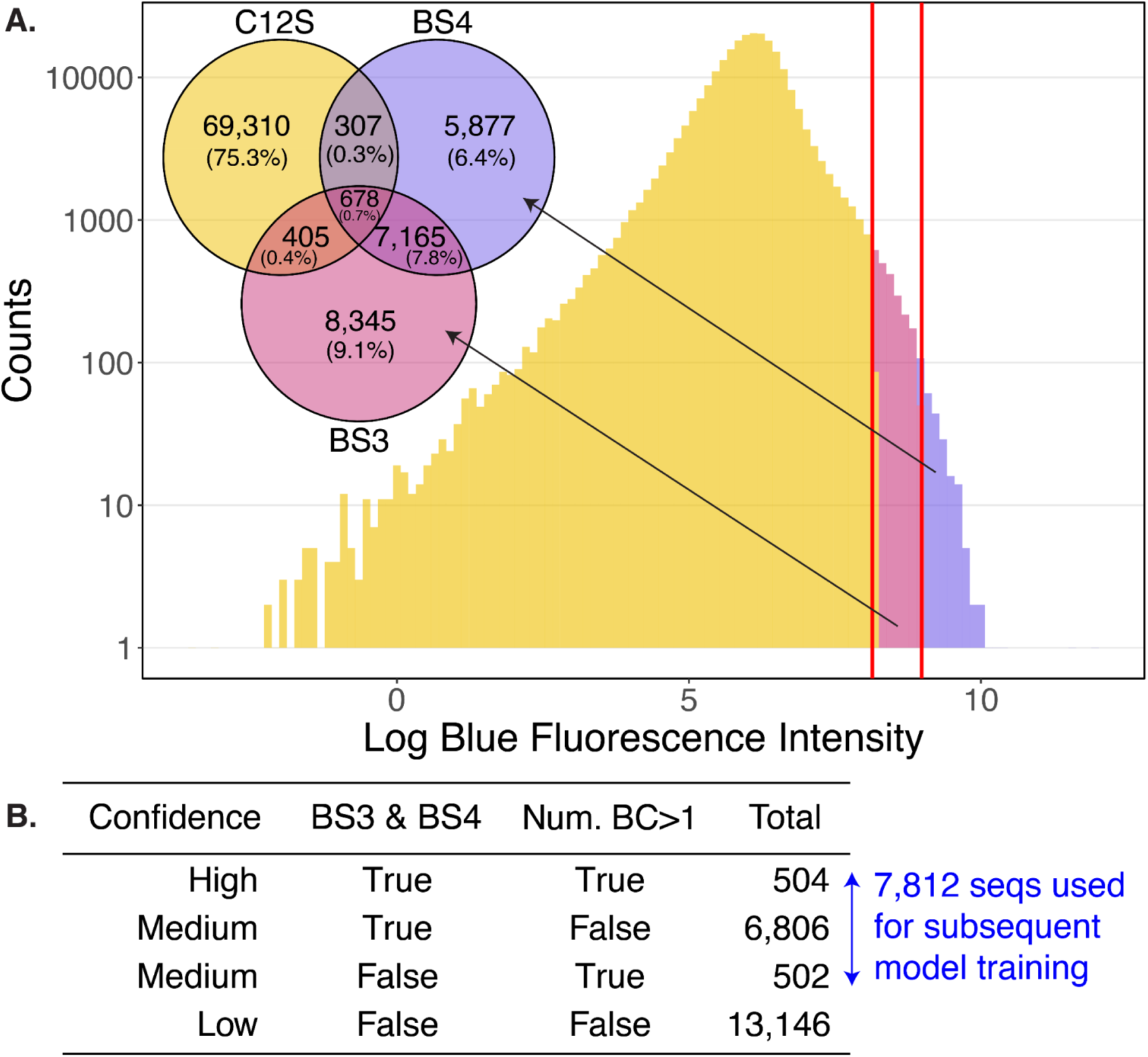
Blue FACS Sort of Chimeric Shuffled Fluorescent Proteins. **A.** FACS of the shuffled library (C12S) to isolate blue-emitting variants. The distribution of log blue fluorescence intensity is shown for the shuffled population. Two gates were defined at the upper end of the brightness distribution, corresponding to BS3 (top 1.3% of cells) and BS4 (top 0.13% of cells), as indicated by red vertical lines. PacBio sequencing of sorted populations reveals minimal overlap with the original C12S library (1.1% for BS3 and 1.0% for BS4) and some overlap between BS3 and BS4 (8.5%), demonstrating strong enrichment of rare, highly fluorescent variants from a diverse background. Venn diagram counts represent unique amino acid sequences. **B.** Confidence classification of sorted variants based on barcode support and bin overlap. Sequences were stratified into four confidence levels using two criteria: presence in both BS3 and BS4 and support by multiple independent barcodes (Num. BC > 1). The combined high- and medium-confidence set (7,812 sequences) was used for subsequent model training.

To assess whether sorted sequences were enriched for actual blue fluorescent protein features, we performed protein level alignments using BLASTp. For each variant, we calculated the fraction of alignment hits corresponding to proteins annotated as blue fluorescent (450 - 490 nm) in FPBase. The median fraction of blue aligned hits per variant was 0 for C12S, 0.10 for BS3, and 0.12 for BS4. Statistical comparison using a t-test showed significant differences between C12S and BS3 (p = 6.9E-11) and between C12S and BS4 (p = 2.2E-14), while no significant difference was observed between BS3 and BS4 (p = 0.26), as shown in Fig. S6. This confirms that FACS sorting successfully enriched variants with sequence similarity to known blue fluorescent proteins. We next examined enrichment patterns by comparing read fractions for overlapping sequences between datasets. The Pearson correlation of fractional abundance was 0.41 between C12S and BS3, 0.43 between C12S and BS4, and 0.99 between BS3 and BS4 (Fig. S7). The strong correlation between BS3 and BS4 suggests that both bins represent a highly consistent enrichment of a shared brightness-associated subset, whereas the weaker correlation with C12S reflects the strong selection pressure imposed by the sort.

Because pooled transformation can result in multiple plasmids entering the same cell, hitchhiker variants may co-segregate during sorting, leading to potential false positives^70,71^. To mitigate this issue and to assign confidence levels to each variant, we analyzed barcode multiplicity. On average, C12S contained 1.3 barcodes per amino acid variant, while BS3 and BS4 each contained 1.2 barcodes per variant, as shown in Fig. S8. Variants supported by multiple independent barcodes provide stronger evidence of enrichment and reduce the likelihood of hitchhiking artifacts. Based on the number of barcodes and bin overlap, we classified sorted variants into three confidence levels. The high confidence set consisted of sequences present in both BS3 and BS4 and supported by multiple barcodes, totaling 504 variants. The first medium-confidence set included sequences present in both BS3 and BS4 but supported by a single barcode, totaling 6,806 variants. The second medium-confidence set comprised sequences supported by multiple barcodes but detected in only one bin, totaling 502 variants. Low-confidence variants were those observed in only one bin and supported by a single barcode, totaling 13,146 variants. These categories are summarized in Fig. 3B. Combining high-and medium-confidence groups yielded 7,812 sequences used for downstream model training. The original training pool consisted of 12,395 sequences, but improvements to the NGS pipeline reduced the final high-quality set to 7,812 sequences. This curated dataset represents a high-confidence functionally-enriched subset of the shuffled sequence manifold.

To assess structural plausibility of the enriched sequences, we sampled 620 variants from the training set and generated AlphaFold2 structural predictions. Pairwise structural comparison using TM-score revealed a median score of 0.968, with a range of 0.567 to 0.988 and a standard deviation of 0.051, as shown in Fig. S9A. This indicates the majority of enriched sequences retain a β-barrel fold, despite extensive recombination and sequence divergence. The presence of a small fraction of lower TM-score models likely reflects either partial structural rearrangements or prediction uncertainty for highly novel chimeras. Collectively, these results demonstrate that FACS based selection effectively isolates a structurally coherent and functionally enriched subset of highly diverse chimeric proteins.

### Machine Learning Generates Novel Fluorescent Proteins

To explore whether expanded and functionally enriched training data could enable generative discovery beyond the natural fluorescent protein manifold, we fine-tuned the protein language model ProtGPT2^5^ on the curated high- and medium-confidence blue fluorescent protein dataset.

After fine-tuning, we generated 11,000 de novo sequences, collectively referred to as the ProtGPT2 BFP (large) library. To maximize diversity and avoid redundancy, we constructed a multiple sequence alignment and inferred a phylogenetic tree. We then iteratively pruned closely related sequences, systematically removing the two most similar sequences at each step until a maximally diverse set of 1,518 sequences remained. This reduced library, ProtGPT2 BFP (trimmed), was designed to broadly sample the learned sequence manifold while minimizing clustering. Structural plausibility of the generated sequences was evaluated using AlphaFold2 structural predictions on a subset of 278 sequences. The median TM-score relative to canonical β-barrel folds was 0.449, with a range from 0.142 to 0.973 and a standard deviation of 0.229 (Fig. S9B). This broad distribution of TM-scores was notable. While many sequences were predicted to adopt β-barrel-like folds, a substantial fraction showed lower predicted similarity. Rather than limiting designs based on structural predictions, we chose to experimentally test a wide range of generated sequences. Given our synthesis capacity, we aimed to empirically evaluate whether AlphaFold2 predictions underestimate foldability for highly divergent designs or whether the model occasionally proposes alternative folds. This strategy allowed direct assessment of model-generated novelty without imposing strong structural priors.

The 1,518 ML-generated sequences were assembled using DropSynth in two independent codon versions, generating libraries BML1 and BML2. Six canonical blue fluorescent protein controls, each encoded in triplicate codon variants, were added for validation, resulting in 1,536 unique designs. Successful assembly was confirmed by gel electrophoresis (Fig. S10). To normalize for potential expression differences between variants, we constructed a new expression vector in which mKate2 was cloned in-frame upstream of each candidate sequence using an alpha-helical linker^55^ (Fig. S11). This dual-fluorescent design enabled relative fluorescence comparisons while controlling for expression variability. Nanopore sequencing of BML1 and BML2 recovered 648 designed protein sequences and 40,879 mutant variants. Of the original designs, 303 were represented only by mutant sequences, with no corresponding perfect variant observed. The assembled ML libraries were then subjected to two independent FACS sorts, gating on the upper half of the blue fluorescence distribution. To define functional variants conservatively, we developed a barcode-support score BC_FL_ (multiplicative across key sorted populations). This multiplicative formulation rewards consistent enrichment across both codon contexts and replicate sorts, while penalizing sporadic or single occurrence events. Applying a threshold of BC_FL_ > 6 reduced the dataset to 201 perfect designs and 653 mutants, which all together aligned to 361 unique ProtGPT2-generated designs showing reproducible fluorescence enrichment. This barcode weighted approach minimized false positives arising from transformation hitchhiking or stochastic enrichment.

To visualize the relationship between ML-generated functional variants and natural fluorescent proteins, we projected ESM-2 sequence embeddings into two dimensions using UMAP (Fig. 4A). Many ML-designed sequences occupy regions that do not overlap with FPBase entries, indicating expansion into previously unsampled areas of fluorescent protein sequence space. Because UMAP is a nonlinear visualization method, we additionally performed multidimensional scaling using BLOSUM62 distances (Fig. S12). The first two principal components explained 24.7% of the total variance, and many ML-derived variants were distant from six canonical controls (EBFP1.5, moxBFP, SBFP2, Electra2, mTagBFP2, and mBlueberry2). This analysis supports the conclusion that the generative model proposes sequences that extend beyond known evolutionary clusters.

**Fig. 4.**
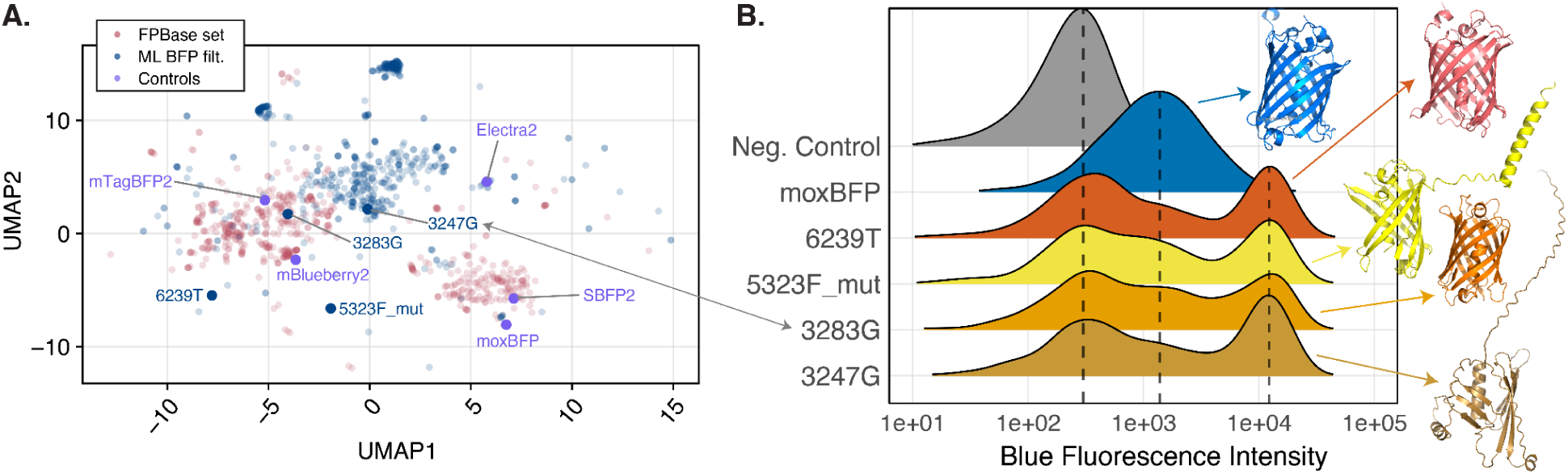
Characterization of ProtGPT2 Generated Blue Fluorescent Proteins. **A.** UMAP projection of ESM-2 embeddings for ML-generated sequences relative to natural fluorescent proteins. Functional ProtGPT2-generated variants (blue) and controls (violet), are visualized alongside FPBase sequences (pink). Many ML-generated variants occupy regions that do not overlap with known natural proteins, indicating expansion of fluorescent protein sequence space beyond the evolutionary manifold. **B.** Experimental validation of dial-out recovered ML designs. Flow cytometry histograms show blue fluorescence intensity distributions of E. coli expressing representative ML-generated variants compared to a positive control (moxBFP) and a negative control. All variants exhibit fluorescence above background. Right panels show AlphaFold3-predicted structures for representative dial-out variants, illustrating diverse predicted structural conformations.

To further validate fluorescence experimentally at the single-variant level, five dial-out PCR recovered proteins, including four perfect designs and one mutant, were individually characterized (Fig. S13; Supplementary Table S1). Flow cytometry (Fig. 4B), plate reader measurements (Fig. S14), and fluorometer measurements (Fig. S15 and Fig. S16) all confirmed detectable blue fluorescence above background controls. Interestingly, AlphaFold3 predictions for several dial-out variants showed incomplete or distorted β-barrel structures (Fig. 4B). This discrepancy suggests either limitations in structure prediction for highly divergent sequences or the possibility that partially altered scaffolds can still support chromophore formation. Previous reports suggest AlphaFold2 and RoseTTAfold can predict chromophore formation^72^. These results demonstrate that fine-tuned generative modeling, when paired with high-diversity experimental training data, can produce functional proteins extending beyond the natural sequence manifold.

### Diversity Expansion Metrics

To systematically quantify how each stage of our pipeline (Fig. 5A) reshaped fluorescent protein sequence space, we applied a complementary set of diversity metrics spanning alignment-based, motif-based, clustering, nearest-neighbor, mosaic structure, and embedding-based analyses. We evaluated the datasets corresponding to each step of the workflow in order: FPBase, C12, C12Shuffled, Blue FACS (Training), ProtGPT2 BFP (large), ProtGPT2 BFP (trimmed), ML BFP synth, and ML BFP filt. As an additional reference for local diversity, we included data from a deep mutational scan of GFP^54^ (Sarkisyan GFP DMS). Together, these analyses provide a multi-scale view of how recombination, functional selection, and generative modeling progressively expand, redistribute, and reconfigure the occupied fluorescent protein manifold.

**Fig. 5.**
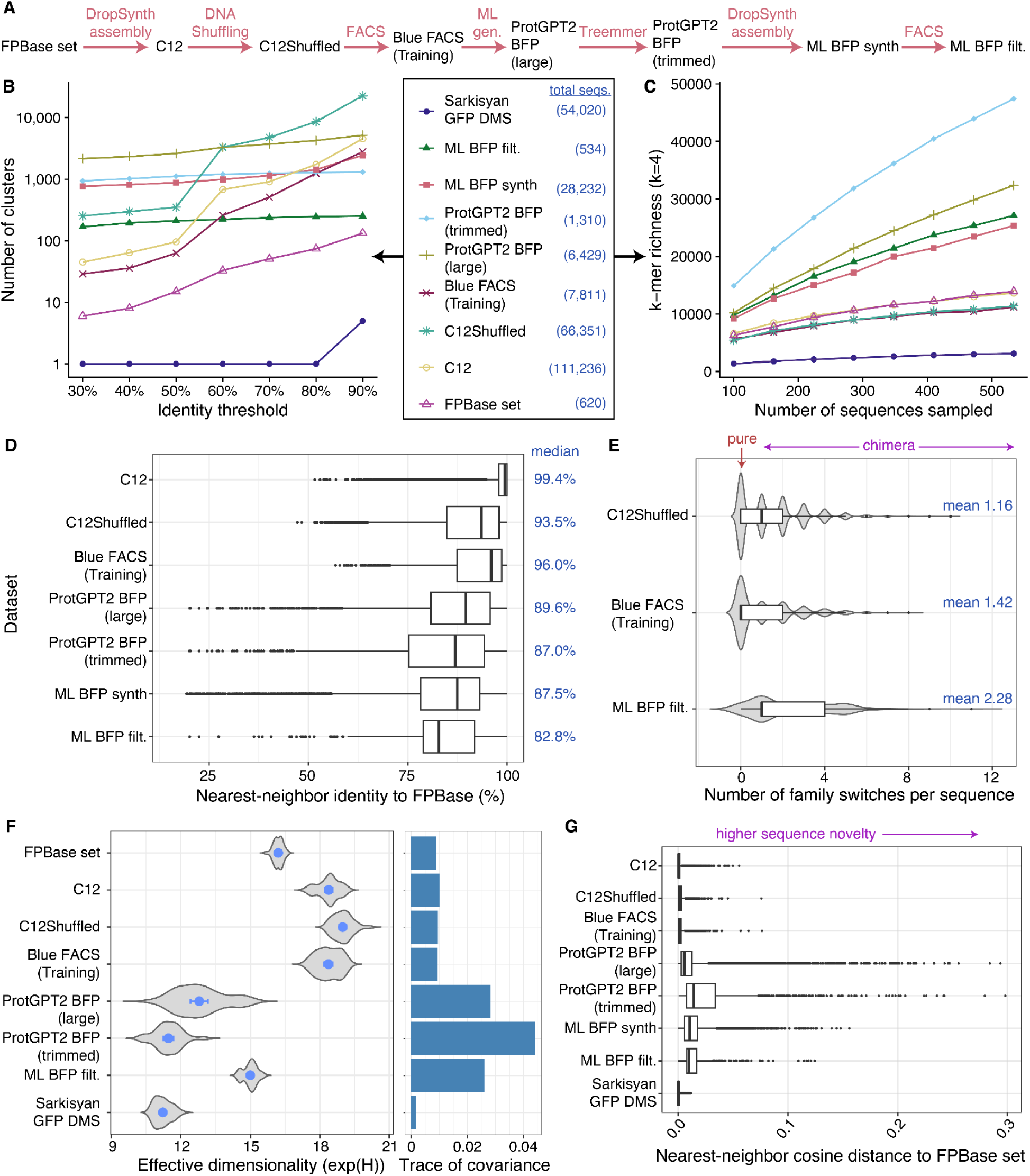
Characterization of Diversity Expansion. **A.** Overview of the datasets in the sequential workflow used to expand and interrogate fluorescent protein sequence space. **B.** Multi-threshold clustering of fluorescent protein libraries. Protein sequences from each dataset were clustered independently using MMseqs2 at amino acid identity thresholds from 30% to 90% with 80% coverage. As expected, cluster counts increase with higher identity thresholds, reflecting finer resolution of sequence differences. Differences in cluster number across datasets illustrate how sequence composition changes at each stage of the pipeline. Because cluster counts are influenced by dataset size and sequencing depth, these results are intended to provide a relative comparison of dispersion patterns rather than absolute measures of diversity. **C.** k-mer diversity. Rarefaction curves for 4-mer richness as a function of the number of sequences sampled, showing mean richness across subsampling replicates. ML-derived libraries accumulate novel 4-mers more rapidly and reach higher richness than parental sets. **D.** Directional nearest neighbor novelty relative to FPBase. Boxplots show the distribution of maximum nearest neighbor percent identities for each dataset when queried against FPBase sequences. Each point represents one sequence in the query dataset, and the distribution summarizes how closely it is represented within the natural FPBase reference. Parental C12 sequences cluster at high identity values, while shuffled and FACS enriched datasets show modest divergence. ProtGPT2 generated libraries and the filtered ML BFP set exhibit broader distributions with lower medians and extended lower tails, including three ML BFP filt variants with nearest neighbor identities below 30%, demonstrating substantial sequence novelty relative to known fluorescent proteins. **E.** Distribution of parental family switches per sequence. Violin and boxplots show the number of transitions between FPBase-defined parental family clusters along each sequence for ML BFP filt, Blue FACS training, and C12Shuffled libraries. ML-derived sequences exhibit a higher frequency and broader distribution of family switches, indicating increased long-range mosaic structure. **F.** Left: Size-controlled effective dimensionality of embedding space. Violin plots show distributions of effective dimensionality exp(H) across repeated subsampling replicates, with points and error bars indicating mean ± 95% confidence interval. Elevated dimensionality reflects exploration along multiple independent axes of variation. Right: The trace of the covariance matrix in PCA space, reflecting total variance across principal components. **G.** Nearest-neighbor cosine distance to FPBase set. Boxplots show distributions of cosine distances from each sequence to its closest FPBase neighbor. Higher values indicate greater embedding novelty relative to natural fluorescent proteins.

### Identity-Based Clustering and Diversity Metrics

To quantify how each stage of library construction reshaped fluorescent protein sequence space, we clustered sequences at identity thresholds from 30% to 90% using MMseqs2 with 80% coverage. Increasing identity thresholds progressively resolve finer sequence distinctions, allowing assessment of both broad evolutionary dispersion and local redundancy. Across all datasets, cluster counts increased monotonically with threshold, but the magnitude and slope of this increase varied considerably. Because absolute cluster counts are influenced by dataset size and sequencing depth, comparisons across libraries are interpreted in relative rather than absolute terms. The curated FPBase set resolved into relatively few clusters across thresholds, consistent with its origin as a collection of naturally evolved β-barrel fluorescent proteins occupying a constrained evolutionary manifold. At 70% identity, FPBase separated into only 51 clusters, indicating substantial shared ancestry despite spectral diversity. DNA shuffling dramatically altered this structure with C12Shuffled showing a pronounced expansion in cluster number across intermediate and high thresholds, resolving into 4,757 clusters at 70% identity. While these values scale with dataset size, the steeper increase in cluster number across thresholds reflects increased sequence dispersion introduced by recombination. This reflects combinatorial recombination of homologous templates, which generates long-range segmental rearrangements rather than incremental mutational drift. The cluster size distribution remained comparatively even, indicating that diversity expansion was not driven by a small number of dominant families but by broad dispersion across sequence space. Functional enrichment by blue FACS selection constrained but did not collapse this diversity. The Blue FACS (Training) dataset exhibited fewer clusters than C12Shuffled at matched thresholds, consistent with selection favoring structurally compatible regions of sequence space, yet it retained substantially greater dispersion than FPBase. Thus, sorting carved out a structured but expansive functional submanifold rather than reverting to a narrow ancestral basin. Generative modeling produced a distinct pattern. The ProtGPT2 BFP (large) library exhibited high cluster counts at low thresholds but plateaued more rapidly at higher identity, suggesting that the model proposes moderately divergent sequences that remain related within broader structural families. Diversity pruning of this set reduced redundancy at lower thresholds while preserving dispersion at higher thresholds, confirming that trimming effectively redistributed sampling across the learned manifold. Notably, the experimentally validated ML BFP filt library retained substantial diversity after functional filtering. Among variants passing the BC_FL_ threshold, clustering at 90% identity yielded 267 clusters, and 224 clusters at 50% identity (Fig. 5B). Although the largest cluster contained 25.0% of sequences, the remaining sequences were distributed across many smaller clusters, indicating that fluorescence selection did not collapse the generative output into a single dominant lineage. Instead, the ML-derived functional proteins occupy multiple distinct regions of sequence space, consistent with expansion beyond natural evolutionary groupings. As expected, the Sarkisyan deep mutational scan control remained largely monolithic across low and intermediate thresholds, fragmenting only at high identity cutoffs. This behavior reflects its origin as a local mutational landscape around a single template rather than a recombination-expanded manifold. Together, these clustering trends demonstrate a progressive reshaping of the fluorescent protein sequence manifold. Natural proteins occupy a compact evolutionary basin, and DNA shuffling broadens this basin through long-range recombination. Functional sorting imposes structural constraint while preserving dispersion. Generative modeling then proposes sequences that populate additional regions within and adjacent to this expanded manifold. Taken together, the relative clustering patterns across thresholds support the central premise of this work: experimentally expanding the training manifold reduces reliance on extrapolation and increases the likelihood that ML-generated designs fall within interpolation regimes of the learned representation.

### k-mer Diversity Analysis

Short amino acid k-mer statistics provide an orthogonal view of diversity that is insensitive to global alignment. Rarefaction analysis demonstrated that observed differences were not solely attributable to sample sequencing depth. When richness was plotted as a function of the number of sequences sampled with replicate subsampling (Fig. 5C), the ML-derived libraries consistently maintained higher 4-mer richness than the C12Shuffled, parental C12, and FPBase datasets, indicating exploration of previously unsampled short-range amino acid motifs. These patterns indicate that generative modeling expands local sequence space in a manner that is robust to normalization for dataset size. Shannon entropy and the corresponding Hill number N1 mirrored these trends (Supplementary Table S2), confirming that increases in richness were not driven solely by rare k-mers, but instead reflected a broader redistribution of motif frequencies. To compare compositional divergence more directly, we computed Jensen-Shannon divergence between each dataset and FPBase as reference (Fig. S17). As expected, divergence values were low for the parental C12, C12Shuffled and the Blue Training set. The ML-derived libraries showed the highest divergence values, particularly at k = 4, reflecting substantial remodeling of local sequence composition. Notably, the ML BFP filt library retained high divergence while remaining functionally enriched, suggesting that short-range motif novelty is compatible with maintenance of the β-barrel fold and blue fluorescence.

### Cross-set coverage and cluster overlap analysis

To determine whether different libraries occupy shared or distinct regions of fluorescent protein sequence space, we pooled all deduplicated, length-filtered sequences and clustered them at 70% amino acid identity. Each resulting cluster represents a coarse-grained region of the β-barrel fluorescent protein manifold, allowing comparison of global occupancy patterns across natural, shuffled, sorted, and ML-derived datasets. Pairwise Jaccard indices based on shared cluster membership reveal a clear hierarchical structure of overlap, as shown in Fig. S18A. Predictably, the parental C12 and C12Shuffled libraries show substantial overlap, reflecting the derivation of shuffled variants from the parental pool. The Blue FACS training set exhibits intermediate overlap with C12Shuffled, consistent with functional enrichment operating within the expanded shuffled manifold rather than reverting toward the original FPBase distribution. In contrast, FPBase displays comparatively low overlap with synthetic and ML-derived libraries, indicating that recombination and generative modeling shift sequence occupancy beyond natural evolutionary clusters. The Sarkisyan GFP DMS remains largely isolated from other sets, consistent with its origin as a single-template local mutational landscape. Among ML-derived libraries, ProtGPT2 BFP (large) and ProtGPT2 BFP (trimmed) overlap strongly with one another, while ML BFP synth and ML BFP filt retain partial overlap with their generative parents, but also occupy distinct clusters following synthesis and functional filtering. These patterns indicate that each stage of the pipeline progressively reconfigures how sequence space is occupied rather than simply resampling pre-existing regions.

The distribution of dataset representation per cluster further clarifies this structure. The majority of clusters are represented by a single dataset, whereas clusters shared broadly across datasets are comparatively rare (Fig. S18B). This predominance of single-dataset clusters demonstrates that most coarse-grained regions of sequence space are uniquely populated rather than universally shared, confirming that shuffling and generative modeling contribute nonredundant coverage. Examination of the 50 largest clusters at 70% identity shows that dominant clusters are frequently populated by C12 and C12Shuffled, with contributions from Blue FACS and ML-derived sequences (Fig. S18C), while FPBase rarely dominates large clusters. ML-derived functional sequences are distributed across multiple large clusters rather than collapsing into a single dominant lineage, indicating that generative design combined with selection does not converge onto a narrow ancestral solution. Together, these cross-set coverage analyses show that DNA shuffling expands the fluorescent protein manifold into previously unoccupied recombinational space, functional sorting reshapes but does not collapse this expansion, and generative modeling redistributes occupancy across both shared and novel clusters. The modest overlap between FPBase and ML-derived libraries and the prevalence of single-dataset clusters indicate that a substantial fraction of functional ML-generated proteins reside in coarse-grained regions not represented in natural datasets.

### Nearest-neighbor cross-coverage analysis

Nearest neighbor cross coverage analysis provided a directional view of how completely each library is represented within FPBase and how far individual sequences extend beyond the natural reference manifold. For each ordered dataset pair, we identified the closest sequence in the reference set and recorded the maximum percent identity, thereby generating full distributions of nearest neighbor identities rather than single summary values. As expected, when FPBase was used as the reference, the parental C12 library exhibited the highest median identity, with most sequences tightly clustered near the upper end of the identity range (Fig. 5D). The C12Shuffled library and the Blue FACS training set showed broader distributions with slightly reduced medians, reflecting recombination-driven divergence while still retaining substantial similarity to natural scaffolds. In contrast, both ProtGPT2 generated libraries and the experimentally filtered ML BFP set displayed systematically lower median nearest neighbor identities to FPBase, with visibly expanded lower tails. This shift indicates that generative modeling followed by functional selection populated regions of sequence space that are not densely sampled by natural fluorescent proteins.

The filtered ML BFP set retained substantial coverage of FPBase clusters while simultaneously exhibiting pronounced novelty. The median nearest neighbor identity to FPBase for ML BFP filt was distinctly lower than that of the parental and shuffled datasets, and its interquartile range extended well below 0.80, indicating that a large fraction of functional ML variants fall outside the dense natural core. Notably, three variants in the final ML BFP filt set exhibited extremely low nearest neighbor identities to FPBase, with values of 20.1%, 22.5%, and 27.4%. These sequences lie near the lower bound of the overall identity spectrum and represent substantial divergence from known natural templates while remaining experimentally validated as fluorescent.

### Chimera-aware mosaic structure analysis

To quantify long-range recombination structure in shuffled and ML-derived fluorescent proteins, we performed a windowed similarity analysis relative to natural FP parents. FPBase sequences were first clustered at 70% sequence identity using MMseqs2, and each cluster was treated as a parental family. This windowed family assignment revealed clear differences in long-range recombination structure across shuffled, experimentally selected, and ML-generated libraries. The shuffled C12S library exhibited a median of one parental family switch per sequence and a mean of 1.42 switches, with 39% of sequences remaining pure single-family constructs (Fig. S19A), as well as an average of 2.15 contributing families per sequence (Fig. 5E). The Blue FACS training set showed more modest mosaicism, with a median of zero switches, a mean of 1.16 switches, 51% pure sequences, and an average of 1.93 contributing families, consistent with partial recombination followed by functional selection that enriches for structurally coherent variants. In contrast, the ML BFP filtered library displayed substantially elevated mosaic structure, with a mean of 2.28 switches per sequence and nearly three parental families contributing on average. The distribution of switches per sequence shows a pronounced rightward shift for ML BFP filt relative to both shuffled and FACS sorted libraries, with a long tail of highly mosaic variants exhibiting numerous family transitions. Consistent with this, the fraction of mosaic sequences is highest in the ML BFP filt set, exceeding 85.3%, compared to 60.7% in C12Shuffled and 48.8% in the Blue training set. These data indicate that the ML model does not simply interpolate within single parental lineages, but frequently assembles sequence segments that most closely resemble distinct natural families along the same polypeptide chain. The positional distribution of recombination boundaries (Fig. S19B) further clarifies this behavior. In the C12Shuffled and Blue libraries, switch densities are enriched in discrete regions along the fluorescent protein scaffold, reflecting recombination hotspots imposed by library design and structural constraints. In contrast, the ML BFP filt sequences exhibit broader and more diffuse boundary distributions spanning much of the sequence length, suggesting that the model combines motifs across multiple structural regions rather than recapitulating predefined junctions. Together with the increased number of contributing families per sequence, these results demonstrate that ML-derived variants occupy a regime of multi-parental mosaicism that exceeds both shuffled and FACS-selected libraries.

### ESM-2 embedding based diversity analysis

Embedding-based analyses using ESM-2^2^ representations revealed that synthetic and ML-generated libraries expand fluorescent protein sequence space along multiple quantitative axes while remaining partially embedded within the natural manifold. UMAP visualization of FPBase and the final ML BFP filt library showed substantial overlap but clear local redistributions, with ML variants populating peripheral regions adjacent to, yet often outside, dense FPBase clusters (Fig. 4A). Labeled examples illustrate that several high-performing ML designs reside at the edges of natural clusters or in sparsely occupied regions, consistent with directional exploration rather than simple recapitulation of known families. Despite these visible shifts, the balanced silhouette score was negative (−0.174), indicating that datasets are not cleanly separable as distinct clusters in full embedding space. Thus, synthetic sequences do not form an isolated cloud, but instead broaden and reshape a shared manifold. Quantitative measures of embedding dispersion support this interpretation. The trace of the covariance matrix in PCA space (Fig. 5F - right), reflecting total variance across principal components, was substantially higher for generative libraries such as ProtGPT2 BFP (trimmed) and ProtGPT2 BFP (large) than for FPBase derived sets, indicating greater global spread. The ML BFP filt library showed smaller but clearly elevated variance relative to natural controls. Similarly, cosine distances to each dataset centroid revealed broader within-dataset dispersion for ML and generative libraries compared to FPBase and DMS variants (Fig. S20), demonstrating that these datasets do not merely shift location but occupy wider regions of embedding space. Effective dimensionality, derived from spectral entropy of PCA eigenvalues, further indicated that diversified and shuffled libraries explore multiple independent axes of variation. In size-controlled subsampling analyses, effective dimensionality remained elevated for C12, C12Shuffled, and Blue Training relative to FPBase, while ML BFP filt and ProtGPT2 libraries showed distinct dimensional signatures consistent with structured exploration rather than random diffusion (Fig. 5F - left). Importantly, these trends persisted after controlling for dataset size, confirming that differences reflect intrinsic geometry rather than sampling depth. Finally, nearest-neighbor analyses relative to FPBase provide a complementary view of novelty. Cosine distance distributions show that ML BFP filt, ML BFP synth, and ProtGPT2 libraries contain substantial fractions of sequences whose nearest FPBase neighbor lies farther away than typical natural variation (Fig. 5G). In contrast, C12 and C12Shuffled remain tightly embedded within the natural envelope, and Sarkisyan GFP DMS variants cluster extremely close to FPBase. Together, these results indicate that ML-guided and generative approaches expand the occupied embedding volume and, in several cases, increase the dimensional complexity of explored space, while still remaining partially continuous with known fluorescent protein structure. The absence of strong cluster separability alongside increased variance and dimensionality supports a view in which synthetic design promotes manifold expansion and interpolation across previously under-sampled directions rather than wholesale departure into disconnected regions.

## Conclusion

This work demonstrates that deliberate expansion of experimentally characterized sequence diversity can reshape the learning problem for protein engineering. By synthesizing and validating hundreds of β-barrel fluorescent proteins, recombining them through DNA shuffling, and enriching rare blue-emitting chimeras by FACS, we generated a large and diverse training set that supports interpolation across an expanded functional manifold. Fine-tuning ProtGPT2 on this dataset enabled generation of thousands of de novo sequences and experimental recovery of fluorescent proteins that extend beyond known evolutionary groupings. Quantitative diversity analyses across identity clustering, k-mer composition, nearest-neighbor novelty, mosaic structure, and embedding geometry show that recombination and generative modeling jointly expand accessible sequence space while maintaining function. Together, these results suggest a scalable framework for ML-guided exploration of small protein families where natural diversity is limited, and where higher-fitness optima may lie far from standard templates.

## Acknowledgements

This project has been made possible in part by a grant from the Chan Zuckerberg Initiative DAF, an advised fund of Silicon Valley Community Foundation. We thank the University of Oregon GC3F core staff for assistance with NGS sequencing. We also thank Jonathan Dorogin, the OHSU Flow Cytometry Core staff, and Cassandra Gonzalez for flow cytometry guidance. A.B. acknowledges support from the University of Oregon NSF NRT-URoL: Molecular Probes and Sensors for Complex Environments and the Giustina Knight Campus Fellowship.

## Data and Materials Availability

Raw PacBio and Nanopore reads of assembled and shuffled gene libraries and FACS sorted variants were submitted to the NCBI Sequence Read Archive under BioProject accession PRJNA1273454 (http://www.ncbi.nlm.nih.gov/bioproject/1282861). Two codon-optimized fluorescent protein parent libraries generated in this study are available to the research community via Addgene as pooled libraries: codon version 1 (C1P), Addgene catalog #245482, and codon version 2 (C2P), Addgene catalog #245483. For each library, pooled plasmid DNA was prepared as a midiprep-scale stock (∼100ug) to support downstream cloning, screening, or sequencing applications. Library sequence files, cloning details, primer information, and complete instructions for reproducing the sequencing and analysis pipeline are available via Addgene with additional scripts on GitHub (https://github.com/PlesaLab/Fluorescent_Protein_NGS_pipeline and https://github.com/PlesaLab/DropSynth_BC_Mapping). NGS mapping files (linking barcodes and gene variants), alignments, and potential dial-out PCR primers for the two parental libraries (C1P and C2P), DNA, and protein reference files are available on FigShare (https://doi.org/10.6084/m9.figshare.30585419). The library plasmid backbone is also available via Addgene catalog #248345, expressing moxBFP that is excisable with NdeI/KpnI restriction sites. Together, these resources enable direct reuse of the physical libraries as well as full reproducibility of the computational analyses described in this work.

## Supplementary Data

Supplementary Information - Methods (Identity-Based Clustering and Diversity Metrics Analysis, Cross-set Coverage and Cluster Overlap Analysis, Nearest-Neighbor Cross-Coverage Analysis, Chimera Aware Mosaic Structure Analysis, EMS-2 Embedding-Based Diversity Analysis), Figures, and Tables.

Blue_FACS_Training.csv/.fasta - All 20,958 sequences in the Blue FACS Training dataset with confidence level, reads counts and barcode counts in BS3, BS4, and C12S.

<u>C1P.genes and C2P.genes</u> - Gene sequences for the two parental FPBase codon libraries. <u>BML1_BML2_proteins_withM.fasta</u> - ML generated protein designs and controls.

BML1_genes_withATG.fasta and BML2_genes_withATG.fasta - Gene sequences for the two parental FPBase codon libraries.

<u>ML_BFP_filt.csv/.fasta</u> - All 854 fluorescent sequences sorted from BML1 and BML2. <u>FPBase_Parents_proteins_noM.fasta</u> - Parental protein sequences from FPBase.

## Methods

### Design and DropSynth Assembly of the Parents Library (C1P and C2P)

Protein sequences were downloaded from FPBase in February 2019. Sequences were length filtered to below 268 a.a to ensure they could be assembled on 5x 300mer oligos. DropSynth oligos were designed using scripts available online (https://github.com/PlesaLab/ds-oligo-design-docker). The 621 genes were split into one library of 384 genes and a second library of 237 genes. Two different codon versions were generated for each gene, for a total of four DropSynth libraries with 1,242 unique genes in total. Oligos were ordered as a 300mer pool from Twist Bioscience. The pool was rehydrated with 100 uL TE and 1 uL of a 1/10 dilution was used to amplify each subpool with 21 cycles (determined by qPCR). Then 1 uL of 25 pg/uL subpool dilution and 18 cycles were used to bulk PCR amplify each library with biotinylated primers, producing 3.7 ug to 4.8 ug of DNA. An nt.BspQI digestion and streptavidin bead reverse-purification was used to remove the short FWD subpool primer section and expose the microbead barcodes as ssDNA, producing 1.7 ug to 3.3 ug of processed DNA. Each library was captured and ligated onto 20 uL of standard 384x barcoded DropSynth beads using 1.3 ug of DNA and 4 uL of Taq ligase in a 100 uL reaction. Beads were washed thoroughly and resuspended in 100 uL Qiagen Elution Buffer. A standard DropSynth assembly reaction was run using 40 uL of loaded beads, 50uL KAPA HiFi, 1uL BSA, 7uL BsaI, and 0.5uL of each suppression compatible primer. Emulsions were created and incubated as described previously^42^. Emulsions were broken with chloroform and purified products were run on a 2% gel for size selection (Fig. S1) of 700 bp to 950 bp range. Extracted amplicons were column cleaned and eluted in 15 uL. Suppression PCR was run using Q5 and 1 uL of size selected product, producing around 1 ug to 4.5 ug of DNA for each library using 4x 50 uL reactions.

### Cloning of Parent Libraries

The vector backbone (pEVBC1) was produced as described previously using round-the-horn PCR^41^. A total of 3.8 ug of vector backbone was digested with 15 U of NdeI and KpnI-HF in the presence of rSAP. For each library 2 ug of DNA was digested with 15 U of NdeI and KpnI-HF. T4 DNA ligation reactions were incubated at 16C overnight with 150 ng of digested vector and 180 ng of digested genes and column cleaned into 8 uL. DH5 NEB Electro Competent Cells were transformed with 1 uL of ligation product. The resulting plates were scraped and used for miniprep or glycerol stocks. DNA from Library 3 (384 genes) and Library 5 (237 genes) were combined together to form the Codon 1 Parents (C1P) library. DNA from Library 4 (384 genes) and Library 5 (237 genes) were combined together to form the Codon 2 Parents (C2P) library. From the glycerol stocks, pooled plasmid DNA from the C1P and C2P libraries was prepared at midiprep scale and deposited in the Addgene repository to enable widespread use by the research community.

### Parents Library Sequencing (PacBio Sequel II) and Analysis

Each of the two codon parent libraries (C1P and C2P) was PCR amplified with Ox_01 forward and reverse primers for C1P, and Ox_02 forward and reverse for C2P (Supplementary Table S3) using Q5 DNA polymerase, 10 ng of template (miniprepped plasmid), in a 50uL reaction for 14-15 cycles. The two libraries were mixed into a single 40 µL sample containing 35 ng/µL of library mixture (20 ul of 24 ng/ul C1 and 20 ul of 46 ng/ul C2) and submitted to the UOregon GC3F core for sequencing on a PacBio Sequel II instrument producing 2.17 million raw CCS reads. Lima was used to demultiplex CCS resulting in 599,594 for C1P library and 1,570,905 reads for C2P library. A custom python script was used to first identify the constant regions flanking the barcode (TGGCTGCGGAAC-20N-GCACGACGTCAG) allowing up to 3 mismatches. The variable region was extracted from each read by scanning for the presence of the NdeI (CATATG) site at the start codon and an end motif (TAAGGTACCTAAGTG) with a stop codon, KpnI cloning site, and some conserved sequence. Barcode counts were collapsed in starcode (1.4) with a distance of 1 using the sphere algorithm. A consensus call was made for each barcode using a simple majority call. All subsequent analysis and plots were carried out in R (4.3.1).

### Gene Shuffling of C1P, C2P, and C12P

Our gene shuffling protocol was derived from Meyer et al^69^, with some small modifications to improve yield. In preparation for DNA shuffling, we made a fresh 10x DNaseI buffer consisting of 1 M Tris-HCl at pH 7.5 and 200 mM MnCl_2_. We then combined the following in the shuffling reaction: 5 uL 10x DNase buffer, 2 μg of linear input DNA, and water for a total volume of 50 uL. The reaction was allowed to equilibrate at 15°C for 5 minutes in a thermal cycler. Subsequently, 1 μL of DNase I, diluted to a concentration of 0.25 U/μl in 1X DNase I buffer immediately before use, was added to the mixture, followed by an incubation period at 15°C for 20 seconds. The reaction was halted by subjecting it to an 80°C incubation for 10 minutes, effectively stopping further enzymatic activity. The DNA fragments were purified from the mixture, with a column cleanup kit (Zymo), obtaining a DNA yield of 30 to 40 ng/ul in 15 uL of water. The incubation time of the DNase I digest was optimized as shown in Fig. S21. To reassemble fragments and create our new chimeric genes, we tried two different PCR mixes and subjected them to a long, progressive hybridization^73^ in the thermocycler. Specifically, we made 100 uL PCR reactions of Phusion Hifi and Q5 Hifi PCR master mixes each with 200 ng of the fragmented library as template and the following cycling conditions: 94°C for 2 minutes; 35 cycles of (annealing temps: 94°C for 30 seconds, 65°C for 90 seconds, 62°C for 90 seconds, 59°C for 90 seconds, 56°C for 90 seconds, 53°C for 90 seconds, 50°C for 90 seconds, 47°C for 90 seconds, 44°C for 90 seconds, 41°C for 90 seconds, and an extension of 68°C for 90 seconds per kb); a final extension of 68°C for 2 minutes per kb. We then used Zymo PCR cleanup columns on the reactions, and obtained 150 ng/ul for Q5, and 60 ng/ul for Phusion in 10 uL elutions. From these elutions, we Q5 PCR-amplified the chimeric genes with skpp504 F/R primer set for 20 cycles. These were column cleaned and run on a gel, extracting the region where the desired fragment should be, as shown in Fig. S22. After gel extraction, the yield was fairly low (∼600 ng total for each), so we re-amplified the fragments with the same Q5 PCR parameters described above, then column cleaned the product and ran them on the gel. We proceeded with cloning the sample into our backbone (as described earlier) and using sequence verification to assess successful reassembly. Briefly, we transformed the ligated library into DH5 NEB Electro Competent Cells. A mixture of 1 ul ligated DNA and 25 ul cells was electroporated, then mixed with 975 mL of SOC recovery media and incubated in the shaker for 1 hour. After the recovery outgrowth, cells were diluted in 1:10 with SOC media. The cells were then spun down to remove the SOC media and resuspended with M9 minimal media before plating. Plates were then incubated at 37°C and RT for over 24 and 48 hours, respectively. After colonies were big enough to pick, we conducted colony PCR (55°C annealing, 1.5 min extension time, 25 cycles total) with pUC19 FWD+REV primers to verify the successful ligation of shuffled genes.

### Cloning the Shuffled Library

The pEVBC1 vector (a derivative of pUC19) was initially PCR amplified with biotinylated primers to produce a 2115 bp amplicon. A total of 1.2 ug of pEVBC1 backbone DNA was digested with 15U of NdeI, 15U of KpnI-HF, and 1U of rSAP in a total volume of 50uL for over one hour. In order to reverse purify the digested vector, we combined the finished digestion reaction with 6uL of the streptavidin beads and 50 uL of B&W buffer, then incubated the mixture for over an hour at 1400 rpm. We Zymo-column cleaned the mixture and obtained 78 ng/uL of digested backbone after eluting with 10uL of elution buffer. The shuffled library was digested similarly with 700 ng of DNA: 15U of NdeI, and 15U of KpnI-HF (and no rSAP). This reaction was cleaned up with a Zymo column kit, and eluted in 10 uL. The Q5 reaction yielded 20 ng/ul, and Phusion reaction yielded 29 ng/uL. We then ligated the digested shuffled library into the digested vector using T4 DNA ligase. Briefly 150 ng of digested vector and 180 ng of shuffled insert (1:3 molar ratio) were incubated with 3,000U of T4 DNA ligase in a reaction of 20uL at 16°C overnight prior to column cleanup and elution in 8uL.

### Fluorescent Library Visualization with the Typhoon Imager

To verify that the shuffled genes were still functional and exhibited fluorescence, a GE Amersham Typhoon Imager was used to scan plates of library-expressing E. coli colonies under different color channels. The imager had four available excitation/emission settings (Cy2 channel 488 nm ex. with a 525/20 nm emission filter, Cy3 channel 532 nm excitation with a 570/20 nm emission, Cy5 channel with 635 nm excitation and 670/30 nm emission, and SypreoRuby with 488 nm excitation and 570/20 nm emission). The plate images were collected with a 50 um resolution and a PMT voltage of 450V. Using ImageJ, we created a pseudocolor composite image by overlaying scans under each channel, as shown in Fig. S3. The fraction of fluorescent colonies was then estimated by manually counting colonies on half of each bacterial plate. Across the counted area, 1,364 colonies were nonfluorescent and 407 were fluorescent, corresponding to ∼23% fluorescent colonies in the library.

### Blue FACS Sort of Shuffled Library (C12S)

Using serial plate dilutions, a glycerol stab of the C12S library was estimated to yield ∼1.36E6 CFUs. To ensure that each square bioassay dish contained >2E6 colonies, 500 µL of SOC medium was mixed with two glycerol stabs of the shuffled library, plated, and incubated overnight. The following day, colonies were harvested by adding several milliliters of liquid medium to each plate and gently scraping colonies into suspension without disrupting the agar surface. Because LB medium exhibited high autofluorescence, cells were pelleted and washed twice with PBS prior to analysis on a Sony SH800 flow cytometer. To enrich the brightest blue fluorescent variants from the C12S shuffled library, mTagBFP2-expressing E. coli was used as a gating control. The upper end of the library’s fluorescence distribution was gated and sorted into “medium blue” (BS3) and “brighter blue” (BS4) subsets. The combined BS3/BS4 gate represented <2% of the total population, with the brighter blue subset approximately an order of magnitude smaller than the medium blue subset (Fig. S23). Sorted cells were immediately plated onto agar and incubated overnight. Colonies were then scraped and used to prepare glycerol stocks, and plasmid DNA was isolated by miniprep for NGS sequencing.

### Shuffled Library Sequencing (PacBio Sequel II) and Analysis

Miniprepped DNA from the shuffled libraries (C1S, C2S, and C12S), as well as from the FACS blue-sorted subpopulations, was amplified using Q5 polymerase with skpp504F and skpp504R primers (74 °C annealing temperature) for 11–15 cycles, as determined by qPCR. PCR products were column-cleaned, yielding 4.3-8.2 µg of DNA in 48 µL.

### Creation of mKate-alpha linker backbone

Our mKate library backbone was derived from the pEVBC1 plasmid^41^ and includes the mKate2 fluorescent protein gene, a downstream alpha-helix linker, NdeI and KpnI restriction sites for library insertion, and a library barcode region (*see* Fig. S11). The original plasmid contained an NdeI site upstream of the mKate2 gene; this site was repositioned to the 3′ end of mKate2, just before the KpnI site. This was achieved by amplifying the 5′ region of the plasmid (excluding the NdeI site) with primers Mkatelinker_plasmid_rev and Mkate_nondeI_for. To complete the modification, we inserted a 188 bp gBlock (linker fragment) downstream of mKate2. This fragment includes a sequence complementary to the 3′ end of mKate2, a flexible hinge, an alpha-helix linker, and both NdeI and KpnI sites. To ensure overlap between the linker and the barcode region, the plasmid was also amplified with a forward primer (Mkatelinker_plasmid_for) that spans the 5′ end of the barcode and overlaps with the linker fragment. Meanwhile, Mkate_nondeI_for and Mkate_nondeI_rev were used to amplify the mKate2 region itself. These three PCR products - the backbone with a terminal NdeI site, the mKate2 insert, and the linker fragment - were assembled using a three-part In-Fusion reaction. The product was transformed into E. coli and colony-picked for sequence validation.

To incorporate barcodes into the plasmid, the assembled mKate-linker plasmid was linearized with KpnI and gel-purified. Around-the-horn PCR was then performed using primers EVBC8_FWD1 and mKateBC_REV1, generating a 2,837 bp product, confirmed by gel electrophoresis. A second low-cycle PCR (5 cycles) with primers EVBC8_FWD2 and mKateBC_REV2 was used to introduce the barcode region, resulting in a 2,887 bp amplicon (not gel-verified due to low concentration). This product was then re-amplified for 16 cycles with 1 ng of template to yield a strong 2,887 bp band, verified on a gel. The final barcoded plasmid was purified using streptavidin bead-based reverse purification and drop dialysis. This barcoded plasmid served as the final expression vector into which the gene library was cloned. To achieve this, library inserts were ligated into the barcoded plasmid vector at a 4:1 insert-to-vector molar ratio using T4 DNA ligase. To ensure sufficient ATP levels, T4 ligase buffer was supplemented with 1 mM ATP (prepared by mixing equal volumes of 10 mM ATP and T4 buffer). Ligation reactions (20 µL) included 0.2 pmol of insert and 0.05 pmol of vector, incubated overnight at 16 °C. A no-insert control was prepared in parallel. Reactions were cleaned using the Monarch PCR cleanup kit and eluted in 7 µL of preheated elution buffer, followed by drop dialysis for 20 minutes. Final DNA concentrations were quantified prior to downstream transformation.

### Design of the ML-Generated Blue FP Library with ProtGPT2

ProtGPT2^5^ is a large language model (LLM) based on the GPT2 architecture with 36 layers and 738 million parameters, pre-trained on the UniRef50 database using self-supervised next-token prediction. Through large-scale sequence training, the model learns an internal representation of protein sequence statistics, including amino acid propensities and global structural tendencies. We fine-tuned ProtGPT2 on a curated dataset of 12,395 unique blue fluorescent protein sequences enriched through FACS sorting and barcode validation. Training used a 90/10 train-validation split and a learning rate of 1E-6. Following model convergence, we generated 11,000 de novo sequences. Of these, 10,000 were generated using single-amino-acid prompts, with each of the 20 canonical amino acids used 500 times, and 1,000 sequences were generated without prompting. Generated sequences were filtered to remove invalid amino acids and sequences longer than 306 amino acids. To maximize diversity, sequences were aligned using Clustal Omega^74^ and a phylogenetic tree was inferred using FastTree^75^. The tree was pruned using Treemmer^76^ by iteratively removing the most similar sequence pairs until 1,518 maximally diverse sequences remained. Six canonical blue fluorescent protein controls (mTagBFP2, EBFP1.5, Electra2, mBlueberry2, moxBFP, SBFP2) were added in triplicate codon variants, yielding 1,536 total sequences. Each sequence was encoded in at least two synonymous codon versions and split into oligonucleotides for DropSynth assembly into two separate libraries, BML1 and BML2.

### Design and DropSynth Assembly of the ML-generated library (BML1 and BML2)

A 300-mer oligonucleotide pool encoding 1,536 machine learning–generated sequences was ordered from Twist Bioscience for DropSynth assembly. Assembly proceeded similarly to the assembly of the parents library described above. We first determined the optimal number of amplification cycles for each oligo subpool using qPCR with biotinylated skpp15_1 and skpp15_2 primers. Library 1 required 19 cycles, and Library 2 required 18 cycles. Bulk amplification was then performed using 7 cycles for Library 1 and 6 cycles for Library 2. Next, Nt.BspQI nicking reactions were carried out for each library using 8,880 ng of DNA for Library 1 and 8,520 ng for Library 2. Nicked DNA was separated from the biotinylated fragment by incubation with streptavidin-coated beads (55 °C, 800 rpm, ≥1 hr), followed by magnetic separation. The supernatant was column-purified and eluted in 60 µL of EB buffer, yielding 2.28 µg (Library 1) and 3.66 µg (Library 2). The processed oligos were then hybridized to barcoded beads for downstream DropSynth assembly. A standard DropSynth assembly reaction was run using 40 uL of loaded beads, 50uL KAPA HiFi, 1uL BSA, 7uL BsaI, and 0.5uL of each suppression compatible primers. Emulsions were created and incubated as described previously. Chloroform was used to break the emulsion, and purified products were run on a 2% gel for size selection between a 700 - 950 bp range. Extracted amplicons were column-cleaned and eluted in 15 uL. Next, suppression PCR was run using Q5 and 1 uL of size-selected product, producing around 1 ug to 4.5 ug of DNA for each library using 4x 50 uL reactions. Suppression PCR products were verified on a gel (Fig. S10). Suppression PCR products were then cloned into the mKate library backbone.

### mKate+/BFP+ Enrichment Sort

Following transformation of the DropSynth-assembled, ML-generated libraries into E. coli Top10 cells, colony PCR revealed that a substantial number of clones contained truncated mKate genes, truncated ML-generated variants, or both. These truncations likely arose during either backbone construction or the DropSynth assembly process, compromising both expression control and variant integrity. To enrich for functional fluorescent variants and ensure the presence of an intact mKate expression cassette, we performed a fluorescence-activated cell sorting (FACS) enrichment on the libraries. (Fig. S23) We prevented bottlenecking the library’s diversity by using minimal plasmid input and maximized transformation scale. Specifically, we transformed 0.1 ng/µL of each library into Top10 cells and plated 50,000–100,000 colonies per plate across 30 plates, yielding over 1.2 million transformants per library. Colonies were scraped, resuspended in LB, pelleted, and washed twice in 1× PBS. Using a BD FACS DIVA sorter, we gated and collected over 6 million cells that were positive for both mKate and BFP fluorescence, using 405 nm and 561 nm lasers and 431/28 and 610/20 filter sets, respectively. mKate and moxBFP were used as fluorescent controls.

### Sequencing read processing and ML-generated variant calling from FACS-enriched libraries

Nanopore reads from each FACS-enriched bin were processed to generate per-barcode consensus nucleotide sequences and translated protein sequences, which were then linked to barcode read-count tables and mapping/annotation outputs (minimap2 SAM and BLASTp tabular files) for downstream analysis in R. Consensus reads were classified as aligned or non-aligned based on primary mappings, and when multiple mappings or BLAST hits existed for a barcode, a single representative assignment was retained using highest MAPQ (mapping) and highest bit score (BLAST), with ties resolved by majority voting where possible and otherwise by selecting a single representative. Perfect variants were called by exact match to the designated variant reference; all other sequences were assigned IDs using the closest matched reference and a SHA256 hash of the observed amino-acid sequence to enable tracking across experiments. To reduce cross-library carryover in codon-diversified libraries sequenced together, barcodes observed in multiple libraries were retained only in the library with higher read support and apparent cross-contaminant nucleotide sequences were filtered based on strong dominance in one library, after which barcode and read counts were recomputed. Protein variants were summarized by collapsing synonymous DNA into a single protein sequence with counts summed across sorted populations and experiments. A barcode-supported fluorescence score, (BC_FL_), was used to define a higher-confidence subset, defined as:

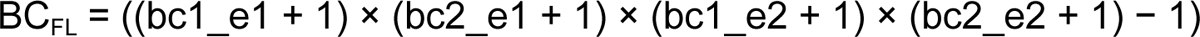

where bc1_e1 and bc2_e1 represent barcode counts in the first FACS sort for codon versions BML1 and BML2, respectively, and bc1_e2 and bc2_e2 represent barcode counts in the second sort. Protein diversity was visualized by MDS on a pairwise distance matrix derived from global pairwise alignments scored with BLOSUM62 (distance = 1 − percent identity).

### ML-generated Fluorescent Protein Characterization

Reads from the mK⁺/BFP⁺ sort were found to cluster into 252 unique ML-generated clusters at 90% identity. Specific BFP variants were then targeted for recovery using dial-out PCR. Variant selection was guided by: I. presence in both codon-optimized libraries; II. a high number of unique barcodes in the library; and III. a length >180 amino acids. Selected variants spanned a wide range of predicted Tm scores, enabling evaluation of whether omitting Tm constraints still yielded functionality. The pEVBC1 plasmid backbone was digested with NdeI/KpnI (37°C, 2 h) and purified by gel extraction. To recover inserts, dial-out primers were designed to amplify target variants from the library, consisting of a forward primer complementary to the start of the BFP variant and a reverse primer complementary to its associated barcode (with the most prevalent barcode cluster selected based on sequencing results, see 6239T-4379K in primer Supplementary Table S3). Using 10 ng of library as template, amplification was performed using touchdown PCR with Q5 reaction mixture under the following conditions: Stage 1, 98°C for 3 min; Stage 2, 98°C for 30 s followed by a 1°C per 30 s rampdown from 76°C to 66°C, then 72°C for 30 s; Stage 3, 95°C for 30 s, 66°C for 30 s, 72°C for 30 s; followed by 72°C for 7 min and a hold at 12°C. PCR products were subsequently cleaned and eluted in 18 µL of elution buffer. Because product concentrations were low, amplification was repeated using a standard PCR protocol (25 cycles; annealing at 66°C). Four replicates per sample were pooled, PCR-cleaned, and eluted in 18 µL of elution buffer. Eluted PCR products were digested with NdeI/KpnI to enable ligation into the pEVBC1 backbone. Digested fragments were analyzed by agarose gel electrophoresis to verify expected sizes (Fig. S13). Because secondary fragments were observed for some samples, gel excision was performed instead of column purification alone. Gel-excised fragments were column-purified, drop-dialyzed for 45 min, and ligated into the digested backbone. Ligation reactions were assembled according to the NEB T4 DNA ligase protocol using 0.06 pmol backbone and 0.12 pmol insert, incubated overnight at 16°C, and heat-inactivated at 80°C for 20 min. Ligation products were purified using the Thermo GeneJET purification kit and eluted in 12 µL of elution buffer, followed by an additional 45 min drop dialysis.

### Flow results of Dial-out Variants

After dial-out variant fragments had been assembled into the pEVBC1 AmpR backbone, several constructs were selected for fluorescence assessment by flow cytometry. E. coli Top10 cells were transformed with the ligation products and cultured overnight at 37°C. Individual colonies were isolated, sequence-verified, and regrown for flow cytometry analysis the following day. Following outgrowth at 37°C, only a small fluorescence-positive subpopulation was detected relative to the negative control, consistent with limited expression and/or impaired folding for most variants. To improve folding, cultures were instead incubated at 18°C for five days, under which fluorescence intensity increased substantially (Fig. 4B). Fluorescence trends were additionally evaluated by plate reader and fluorometer measurements. For these assays, cultures were grown in liquid for three days and then incubated statically at 18°C for one week. Cells were pelleted by centrifugation, washed twice with PBS, and resuspended for measurement. For plate reader assays, 150 µL of each sample was dispensed in triplicate into a clear 96-well plate for OD600 measurement and then transferred to a black flat-bottom 96-well plate for fluorescence detection. Samples were excited at 355 nm, and emission spectra were recorded from 400–700 nm in 2 nm increments on a Synergy Neo2 plate reader. Fluorescence values were averaged across triplicates and normalized to OD600 (Fig. S14). The brightest plate reader sample (2771Q) was subsequently characterized by fluorometry. For fluorometric measurements, 2 mL of each sample was measured for OD600 using a cuvette reader, followed by acquisition of fluorescence spectra on a standard fluorometer (excitation at 355 nm; emission 365–700 nm). Fluorescence intensity was normalized to OD600 and compared against a negative control (E. coli Top10 transformed with pUC19) and a positive control (moxBFP) (Fig. S16).

