## Supplementary Information for "High Diversity Gene Libraries Facilitate Machine Learning Guided Exploration of Fluorescent Protein Sequence Space"

### Supplementary Methods

#### Identity-Based Clustering and Diversity Metrics Analysis

All deduplicated and length-filtered amino acid sequences were clustered independently using MMseqs2 easy-cluster with minimum sequence identity thresholds of 30%, 40%, 50%, 60%, 70%, 80%, and 90%. Clustering required 80% coverage of the shorter sequence with coverage mode 0. For each clustering threshold, the resulting cluster assignment files were parsed to determine the total number of clusters, the total number of unique sequences, cluster size distributions, Shannon entropy H of cluster sizes, effective diversity $N_1 = \exp(H)$, Gini coefficient, largest cluster size, and largest cluster fraction. Shannon entropy and effective diversity quantify how evenly sequences are distributed across clusters, while the Gini coefficient captures dominance of large clusters. This multi-threshold approach provides a scale-dependent view of sequence diversity, distinguishing broad evolutionary separation from fine-grained local variation.

#### Cross-set Coverage and Cluster Overlap Analysis

To quantify the extent to which different sequence libraries occupy overlapping or distinct regions of fluorescent protein sequence space, we performed a cross-set coverage analysis based on sequence clustering. All deduplicated, length filtered protein sequences from every dataset were pooled into a single FASTA file, with sequence identifiers annotated to preserve dataset origin. The combined sequence set was clustered using MMseqs2 at a 70% sequence identity threshold, generating a global set of sequence clusters that represent coarse-grained regions of protein sequence space. For each resulting cluster, we recorded the presence or absence of sequences from each dataset, thereby constructing a cluster-dataset incidence matrix. Using this representation, we computed (i) the fraction of total clusters containing at least one sequence from each dataset, as a measure of global cluster coverage, and (ii) pairwise Jaccard indices between datasets (Fig. S18A), defined as the size of the intersection of clusters occupied by both datasets divided by the size of their union.

Nearest-Neighbor Cross-Coverage Analysis

To quantify asymmetric coverage and novelty between sequence libraries, we performed nearest-neighbor cross-coverage analysis based on pairwise sequence similarity. For each ordered pair of datasets (A,B), we computed the nearest neighbor of every sequence in A within dataset B using MMseqs2 alignments, recording the maximum pairwise percent identity for each query sequence. This produced a distribution of nearest-neighbor identities describing how well sequences in A are represented within B. From these distributions, we computed summary statistics including the median nearest-neighbor identity, 10th and 90th percentiles, and the fraction of sequences in A whose nearest neighbor in B exceeded specified identity thresholds. This analysis was repeated in both directions (A->B and B->A), enabling quantification of both coverage (the extent to which one dataset spans another) and novelty (the fraction of sequences that remain distant from a reference set).

Chimera Aware Mosaic Structure Analysis

To quantify long-range recombination structure in shuffled and ML derived fluorescent proteins, we performed a windowed similarity analysis relative to natural FP parents. FPBase sequences were first clustered at 70% sequence identity using MMseqs2, and each cluster was treated as a parental family. Sequences from C12Shuffled, Blue FACS (Training), and ML-generated libraries were segmented into overlapping windows (30 amino acids, step size 10 amino acids). Each window was searched against the FPBase database using MMseqs2, and the best matching FPBase sequence was assigned to its corresponding parental family. Because alignments were performed against clustered FPBase representatives rather than all individual natural sequences, closely related parental variants were collapsed into single families. As a result, recombination events occurring between highly similar natural homologs are not resolved as distinct switches, and the number of inferred family transitions likely underestimates the true frequency of segmental recombination. For each full-length sequence, the ordered series of family assignments was used to compute the number of family switches along the sequence, the number of distinct parental families contributing segments, and the fraction of sequences composed entirely of a single family.

EMS-2 Embedding-Based Diversity Analysis

Protein sequence diversity was further analyzed using representations from a pretrained protein language model. All sequences were embedded using the ESM-2 model (esm2_t33_650M_UR50D), with per-sequence embeddings obtained by mean pooling over residue representations from the final transformer layer. Embeddings were L2-normalized and analyzed using cosine distance. Within each dataset, embedding-space diversity was quantified by (i) the trace of the covariance matrix computed in principal-component space, (ii) the distribution of cosine distances to the dataset centroid, and (iii) effective dimensionality derived from the spectral entropy and inverse participation ratio of PCA eigenvalues. To assess overlap between datasets, a balanced silhouette score was computed using cosine distance. Cross-dataset coverage was evaluated using nearest-neighbor analysis: for each ordered pair of datasets A->B, the cosine distance from each sequence in A to its nearest neighbor in B was computed. A coverage threshold was defined from the 95th percentile of within-FPBase nearest-neighbor distances, enabling estimation of the fraction of sequences in A that fall within the natural FP embedding envelope of B. Dimensionality reduced visualizations were generated using UMAP for qualitative inspection only. All quantitative conclusions were derived from full-dimensional embedding analyses.

**Supplementary Figures**

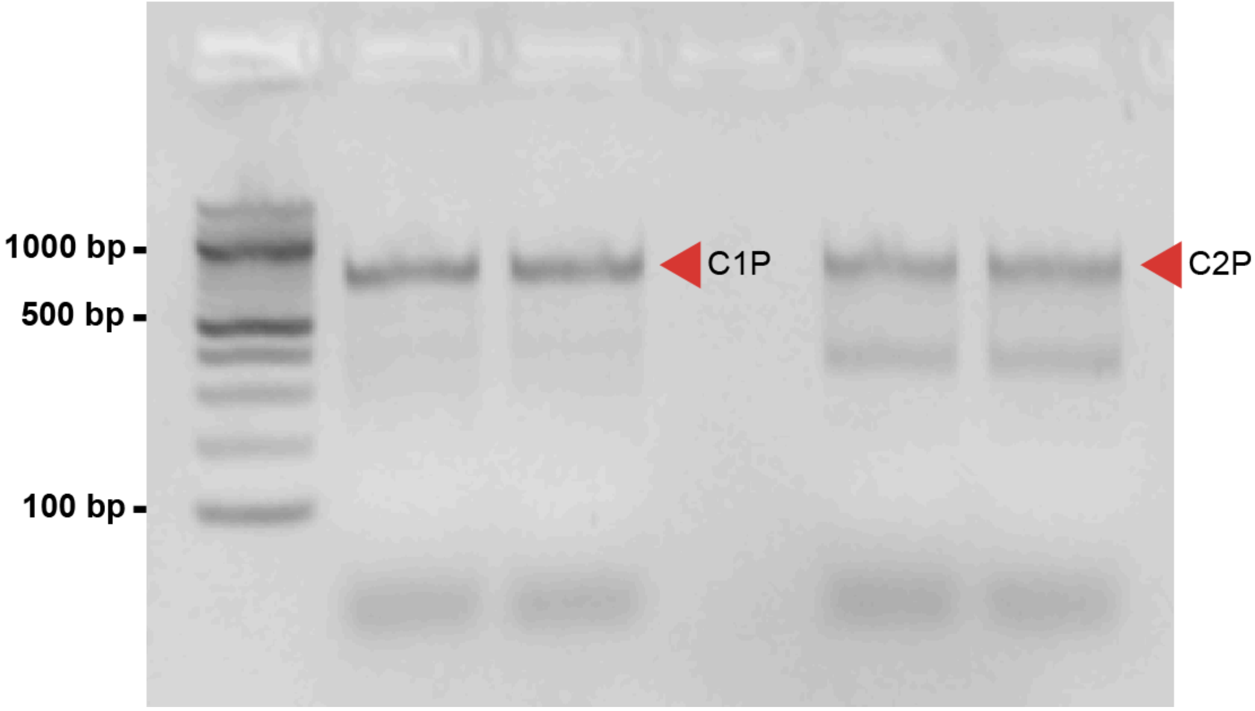

**Fig. S1**. A 2% gel with NEB 100 bp ladder and the two DropSynth assembled libraries C1P and C2P after suppression PCR (loaded in gel as duplicates). Marked bands were gel-extracted prior to cloning.

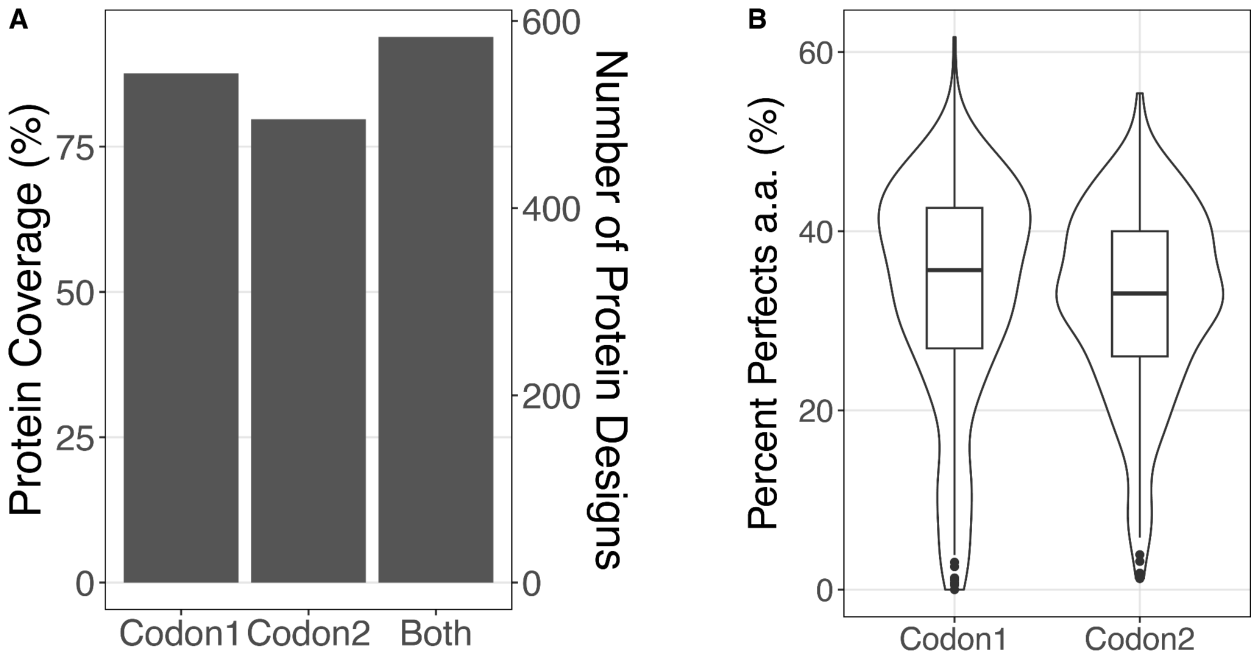

**Fig. S2**. Assembly statistics for parental libraries C1P (codon 1) and C2P (codon 2). **a)** The observed coverage for each individual library and combined across both. A design is counted if we observe at least one perfect assembly in the PacBio sequencing data. **b)** The percentage of perfects observed for each library at the amino acid level (synonymous mutations collapsed) for designs with at least 100 barcodes.

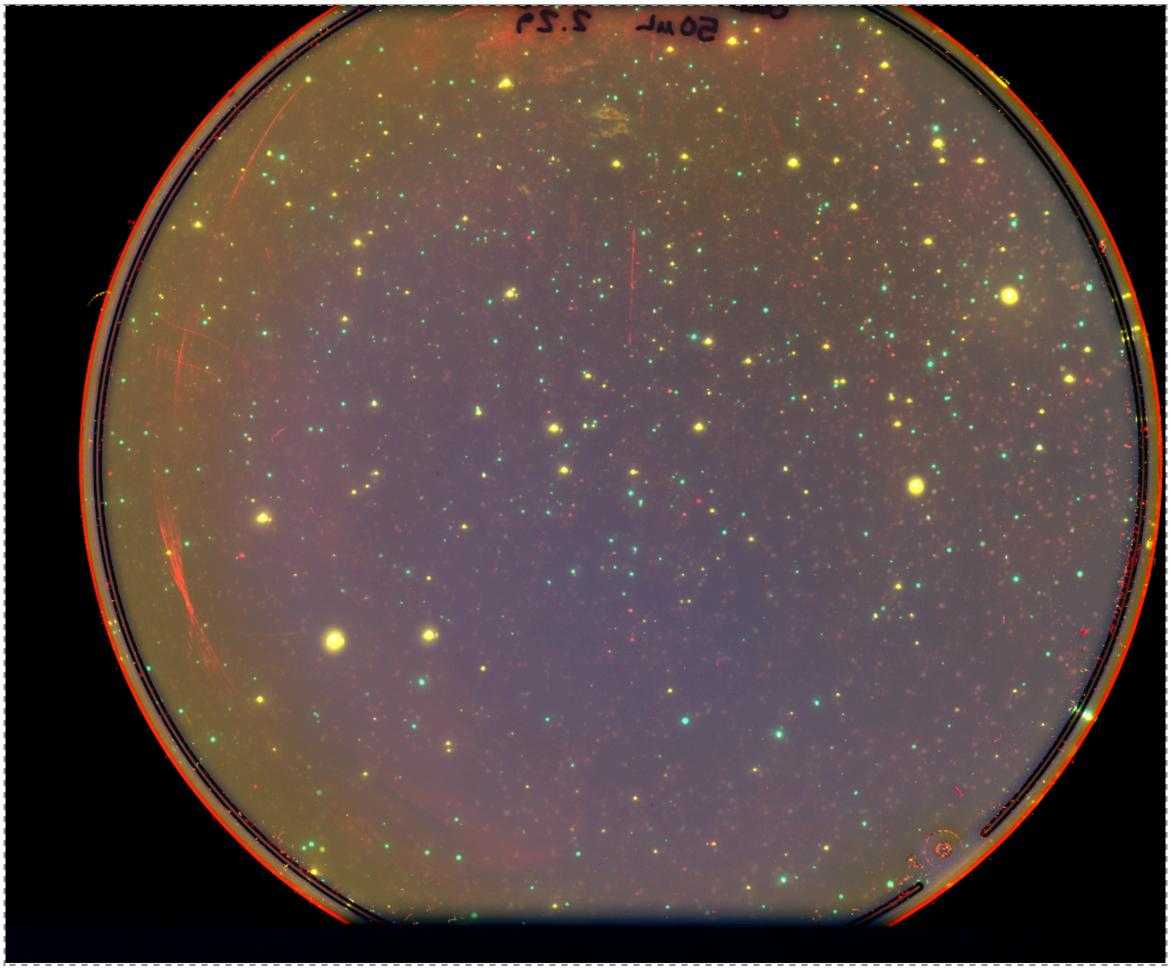

**Fig. S3**. False-color composite image of a shuffled library bacterial colony plate acquired using the Amersham Typhoon scanner. Fluorescent signals from green, yellow, and red channels were overlaid using ImageJ.

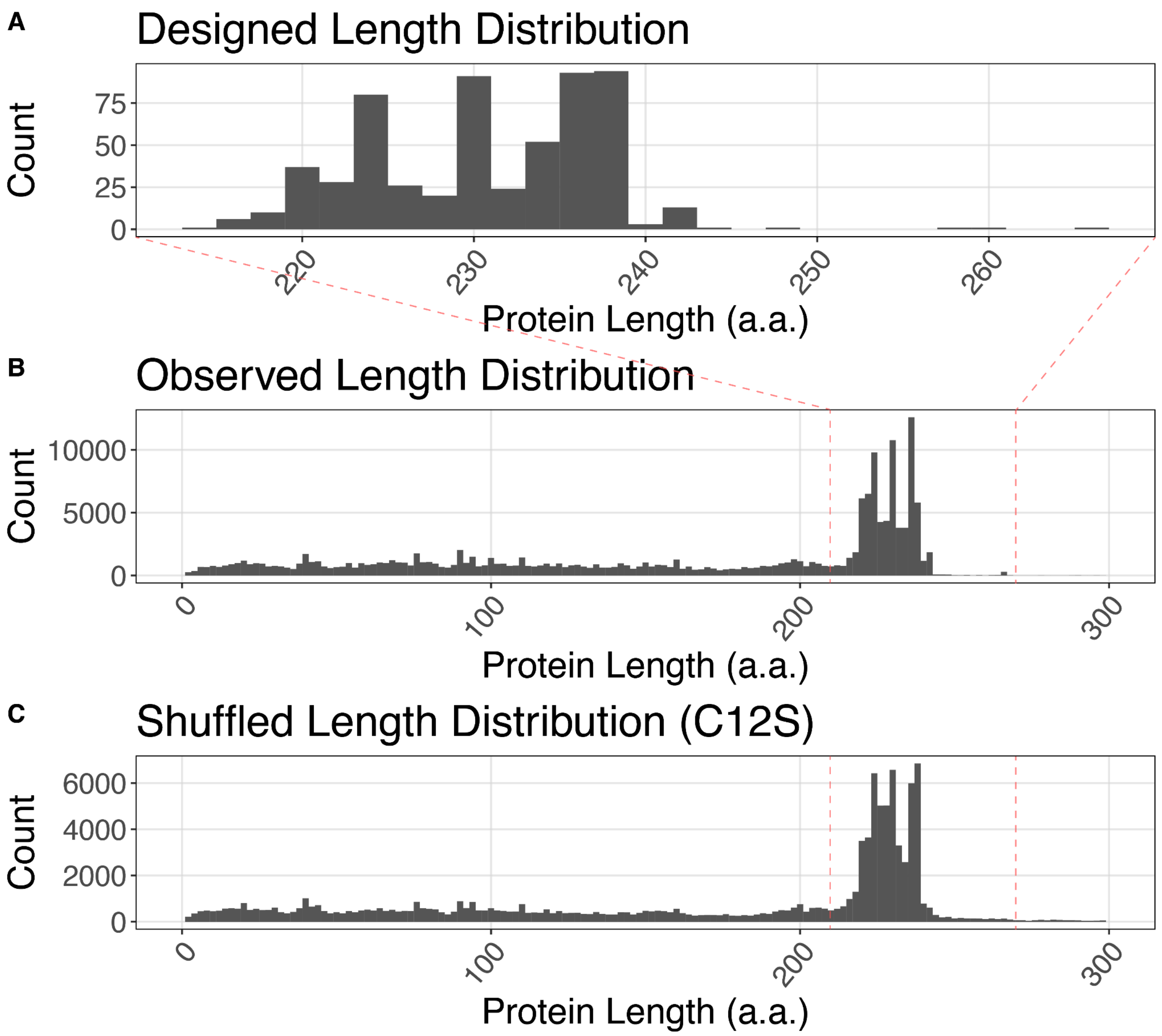

**Fig. S4**. Protein length distributions of **a)** Original 620 FPBase variants. **b)** Observed lengths in parental libraries (C1P, C2P). **c)** After shuffling, in library C12S. Data based on PacBio sequencing.

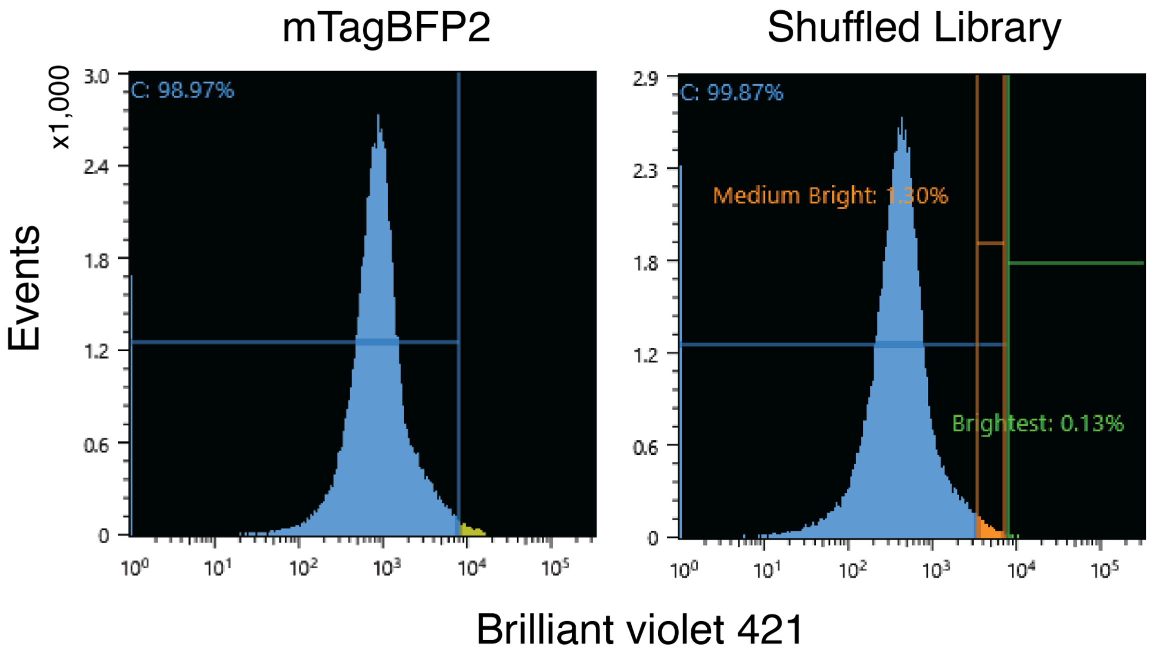

**Fig. S5**. (Left) Flow cytometry histogram of E. coli expressing mTagBFP2 in a pBAD plasmid. A gate was applied to highlight the brighter portion of the fluorescence distribution, used as a reference for blue-positive cells. (Right) Using the blue control gate, a medium blue gate (BS3) was drawn in the shuffled library's distribution to represent the brightest edge of the blue control gate (orange interval gate), and a brighter blue gate (BS4) was drawn to sort anything that was brighter than the entire blue control's distribution (green interval gate).

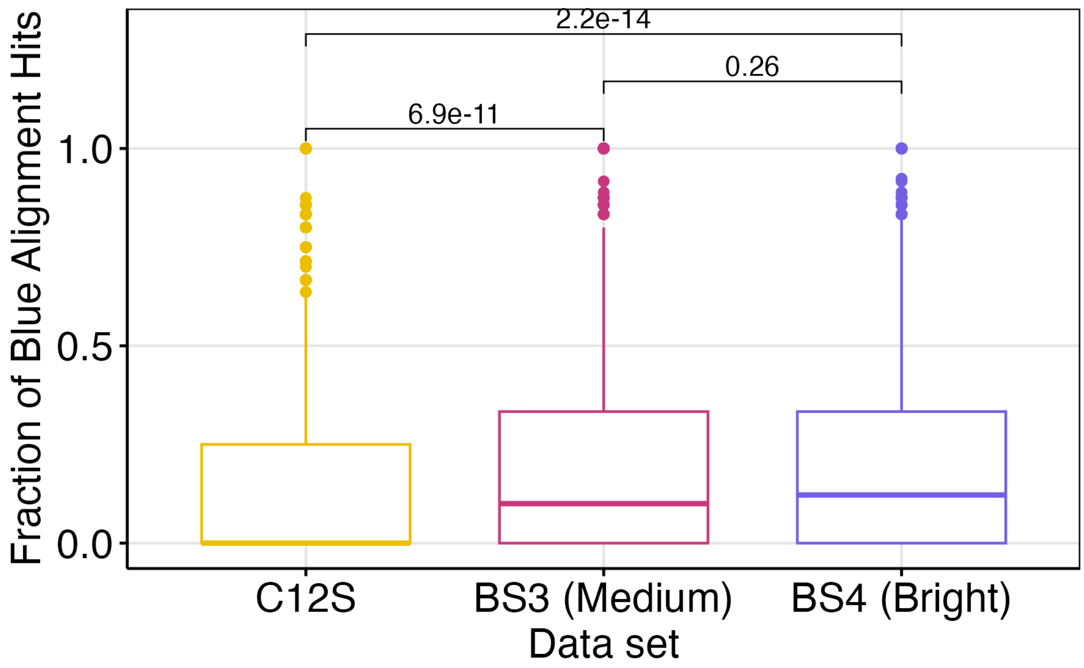

**Fig. S6**. **Fraction of blue alignment hits across shuffled and sorted datasets.** For each unique variant in C12S (unsorted shuffled), BS3 (medium blue), and BS4 (brighter blue), BLASTp hits were annotated using FPBase emission wavelengths and the fraction of hits labeled as blue fluorescent (450-490 nm) was calculated. Blue-hit fractions were significantly higher in BS3 and BS4 than in C12S (t-test; C12S vs BS3 p = 6.9E-11; C12S vs BS4 p = 2.2E-14), with no significant difference between BS3 and BS4 (p = 0.26).

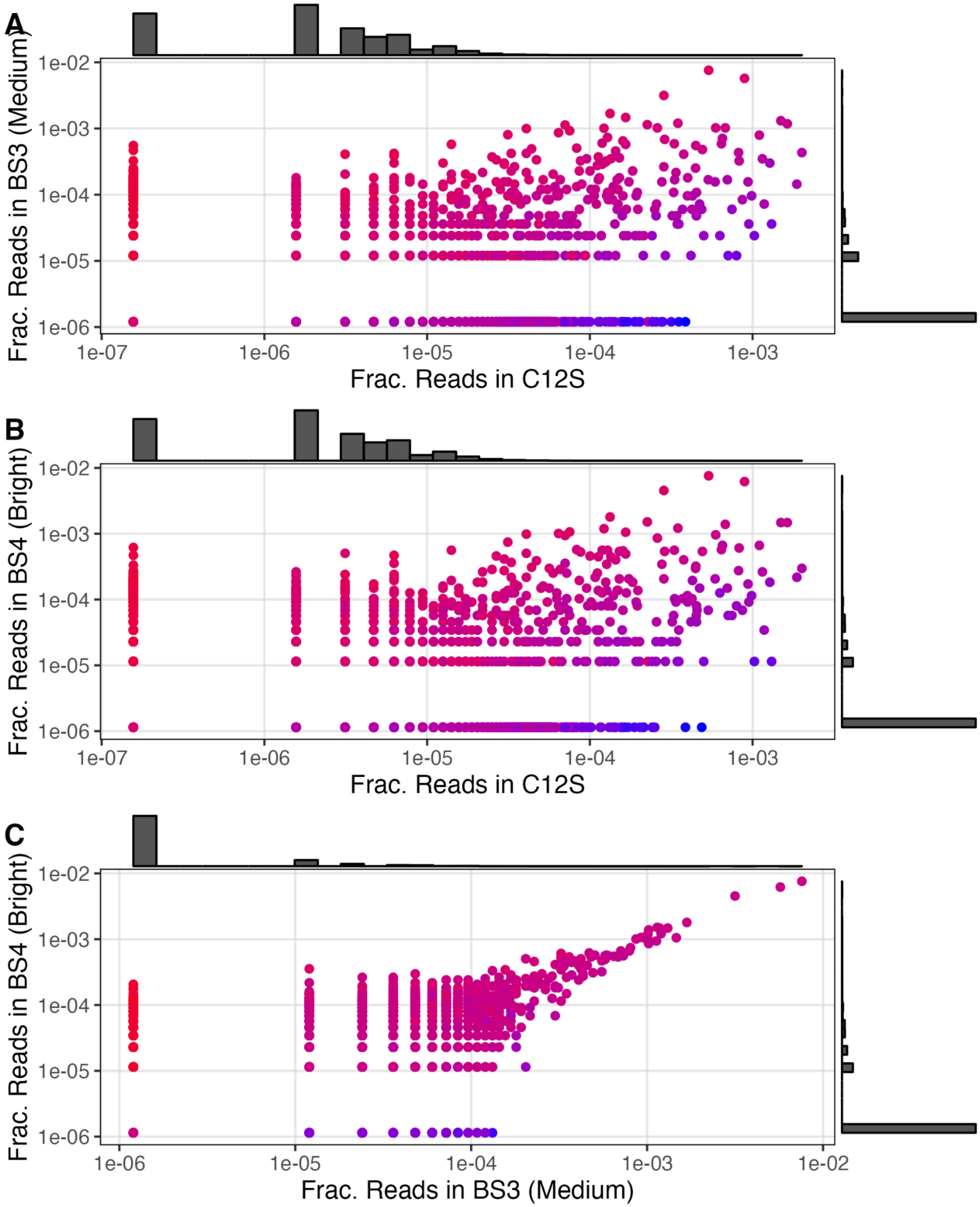

**Fig. S7**. Pairwise comparisons of fractional read abundance across C12S, BS3, and BS4. Shared variants were plotted by their fractional read abundance in each dataset (C12S vs BS3; C12S vs BS4; BS3 vs BS4).

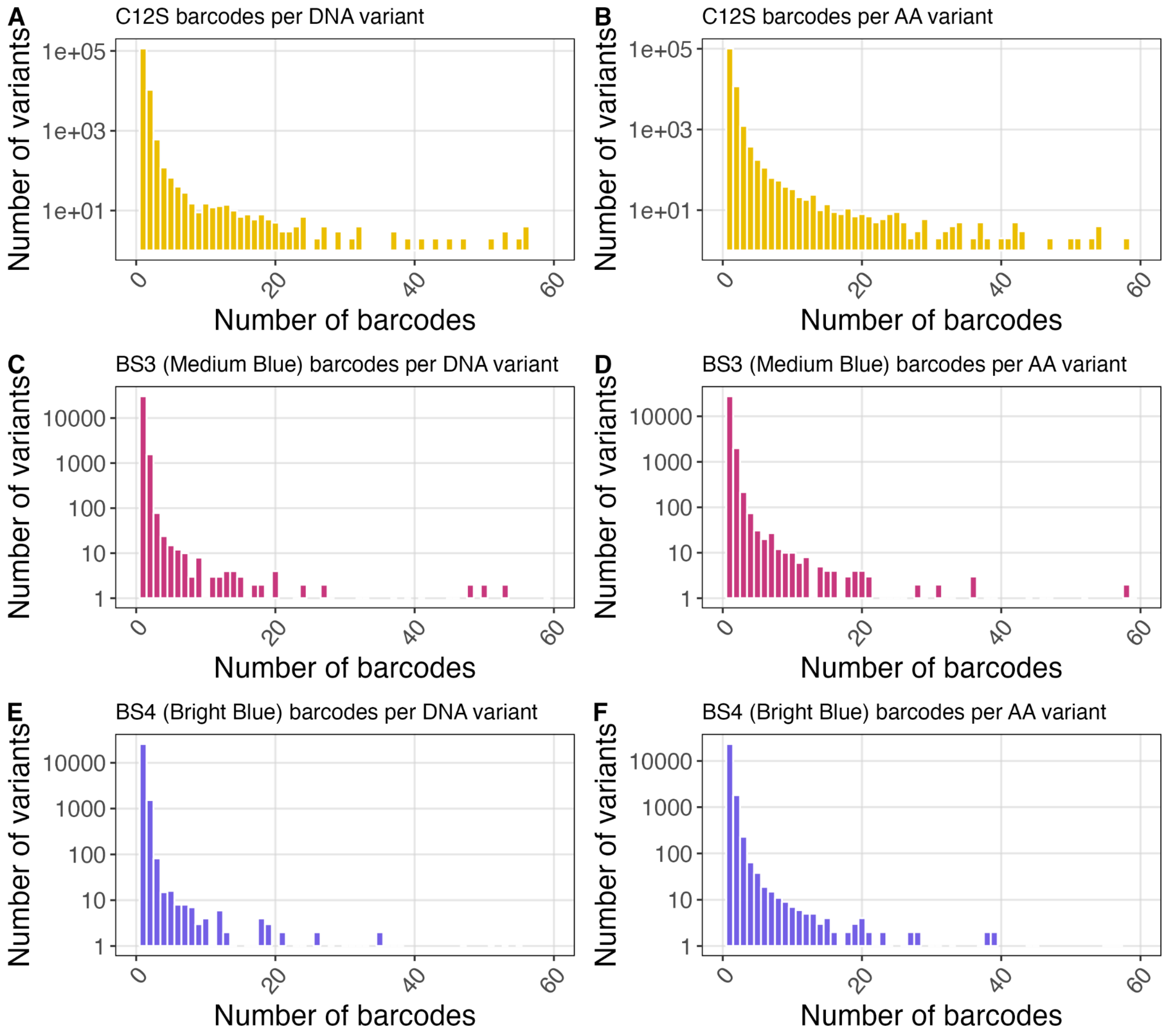

**Fig. S8**. Barcode multiplicity per variant across unsorted and sorted libraries. Distributions of unique barcode counts per DNA variant (left) and per amino-acid variant (right) for C12S, BS3, and BS4, as determined by PacBio sequencing. Barcode multiplicity was used as an indicator of variant confidence and to reduce potential hitchhiking effects from pooled transformations.

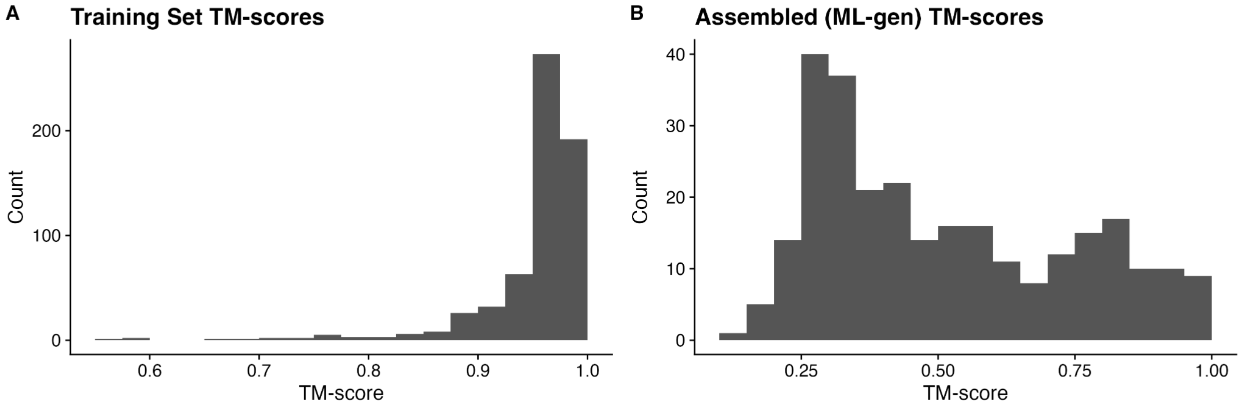

**Fig. S9**. (a). Distribution of TM-scores within the training set relative to various β-barrel fluorescent protein structures listed in FPBase. We observed a median 0.968, range 0.567-0.988, SD 0.051. (b) For a subset of 278 ML-generated sequences, the maximum TM-score was: median 0.449, range 0.142-0.973, SD 0.229.

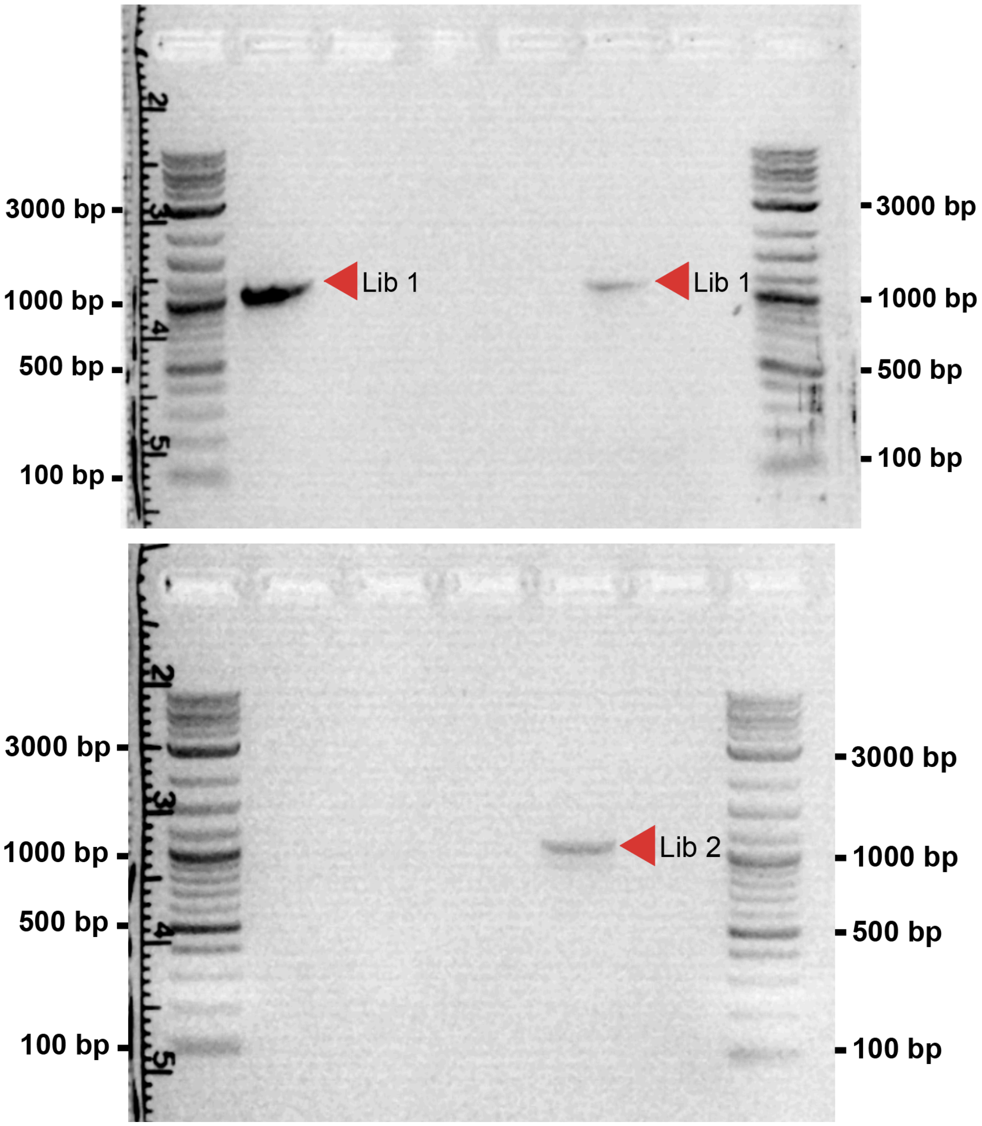

**Fig. S10**. Suppression PCR results of ProtGPT2-generated FP libraries BML1 and BML2, used to amplify products after the blind gel extraction. Products were visualized alongside the NEB 1 kb Plus DNA ladder for size reference. Library bands are ~1120 bp.

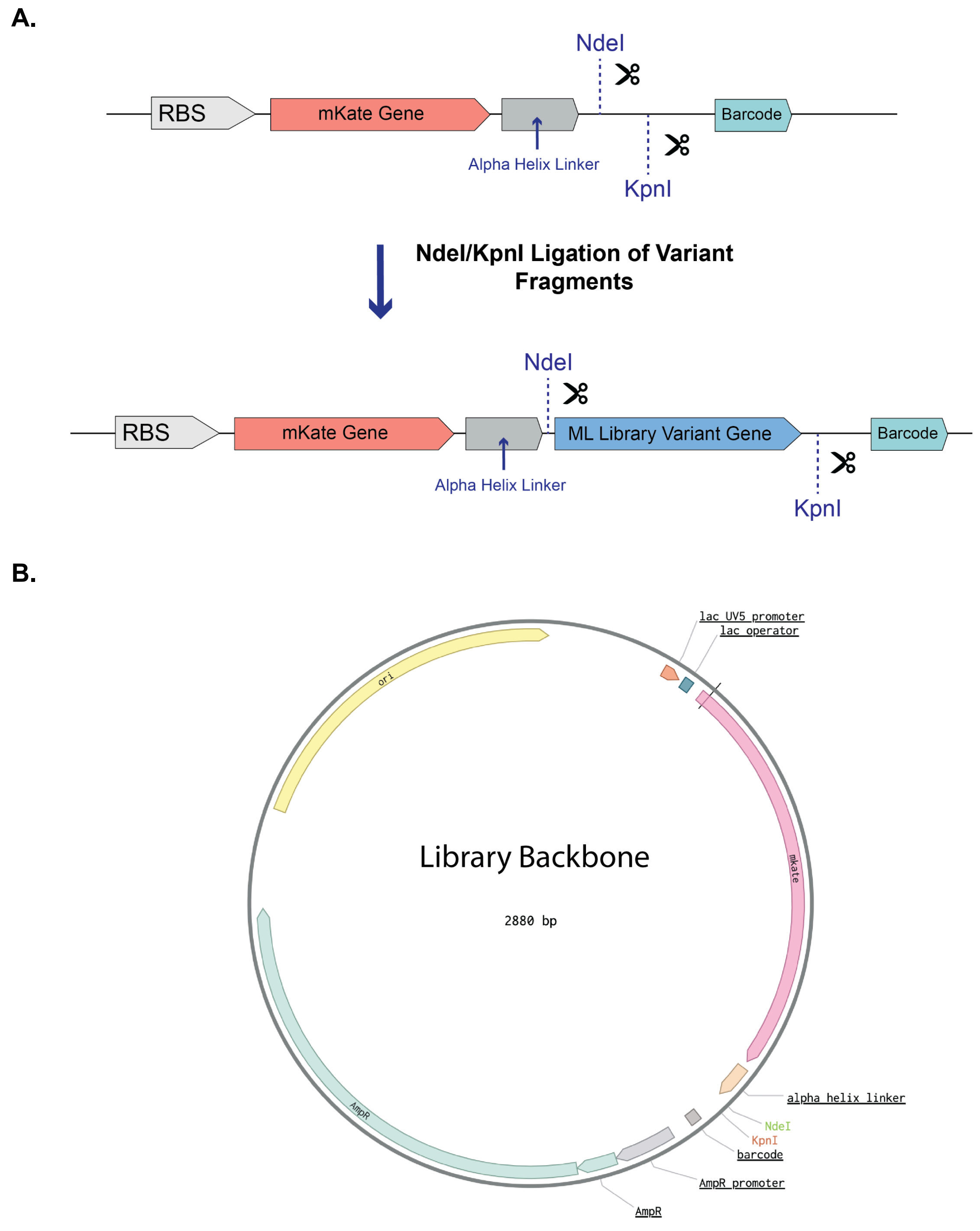

**Fig. S11**. Plasmid maps for library cloning. (a) Expanded view of the library insertion region, showing the NdeI/KpnI cloning site positioned between the α-helix linker and the NGS barcode. (b) Full plasmid backbone map shown without a ligated library insert.

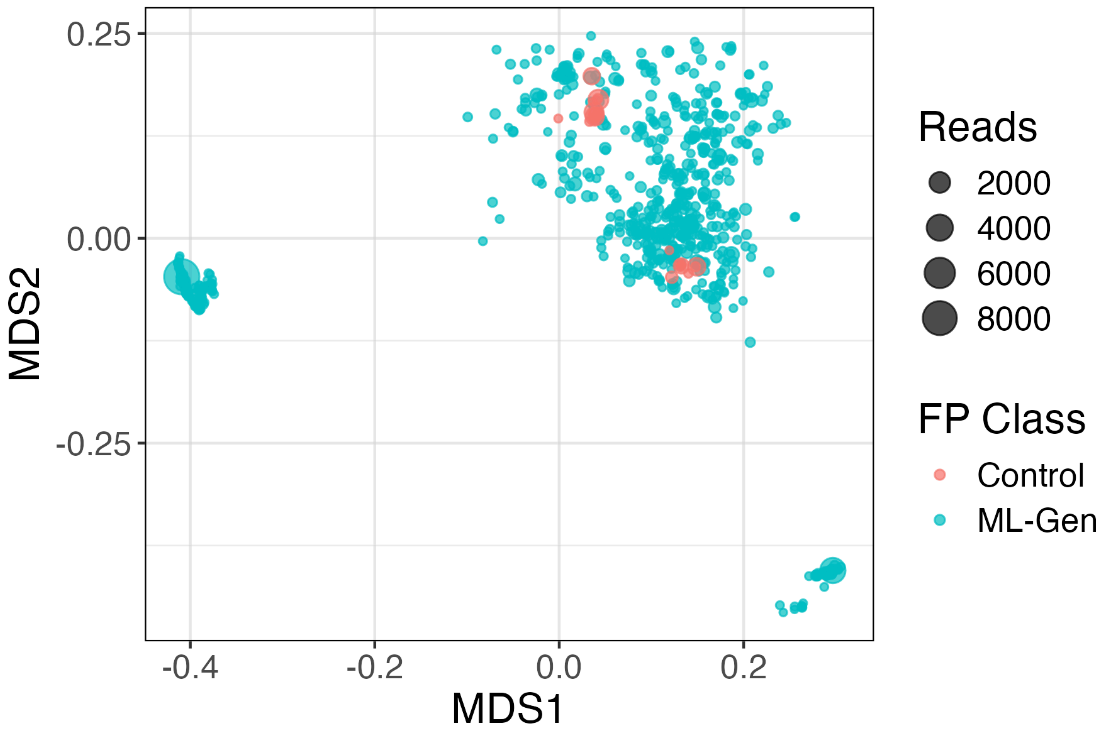

**Fig. S12**. Multidimensional scaling of ML-generated fluorescent proteins. Two-dimensional multidimensional scaling (MDS) projection based on pairwise BLOSUM62 distances between amino acid sequences. Each point represents a unique protein variant, with point size proportional to sequencing read count. ML-generated variants (teal) are shown alongside canonical control fluorescent proteins (red). The distribution indicates that ML-generated sequences occupy a broad region of sequence space, with several clusters positioned away from control proteins, consistent with substantial sequence divergence. Control variants cluster more tightly, reflecting their closer evolutionary relationships. The spread of ML-generated points across MDS space supports the conclusion that generative modeling produces diverse sequences that extend beyond the immediate neighborhood of established blue fluorescent protein templates.

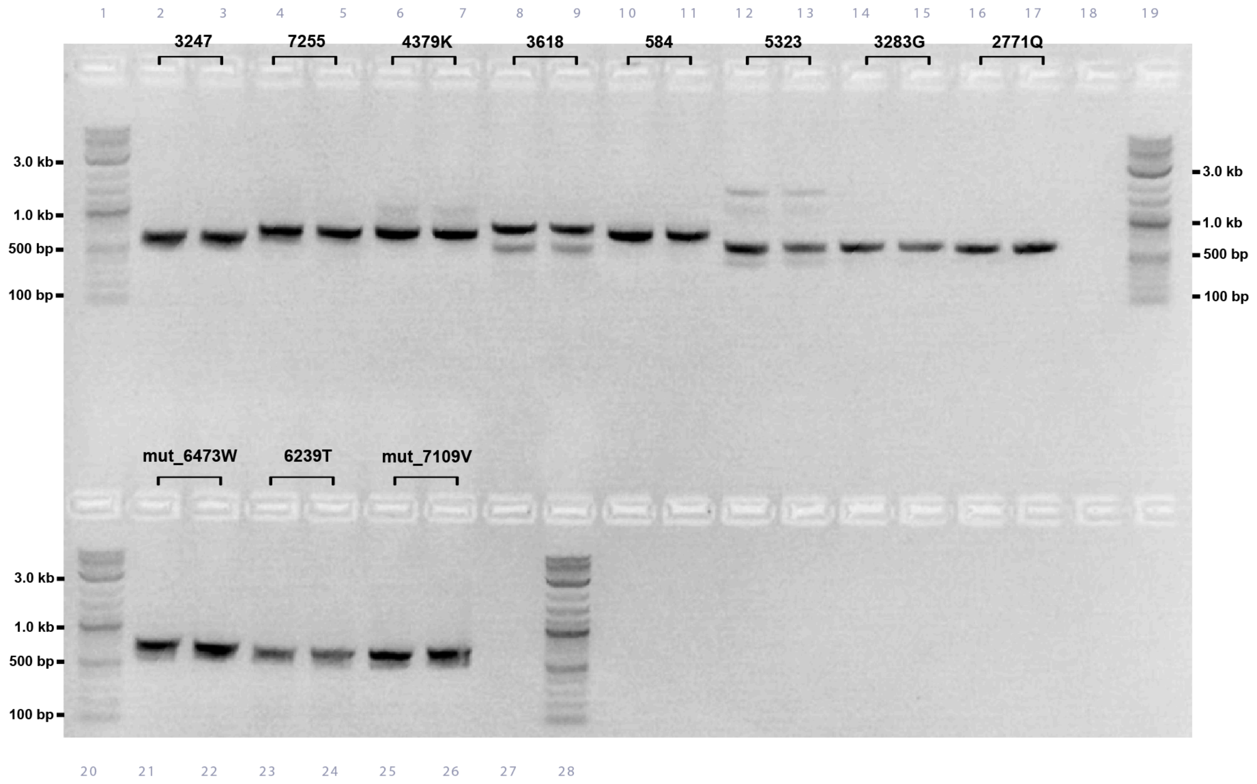

**Fig. S13**. Gel result of variants that were extracted from the ML library via dial-out PCR. Order of samples from left to right (in duplicate): 3247, 7255, 4379K, 3618, 584, 5323, 3283G, 2771Q, mut_6473W, 6239T, mut_7109V. Products were visualized alongside the NEB 1 kb Plus DNA ladder for size reference.

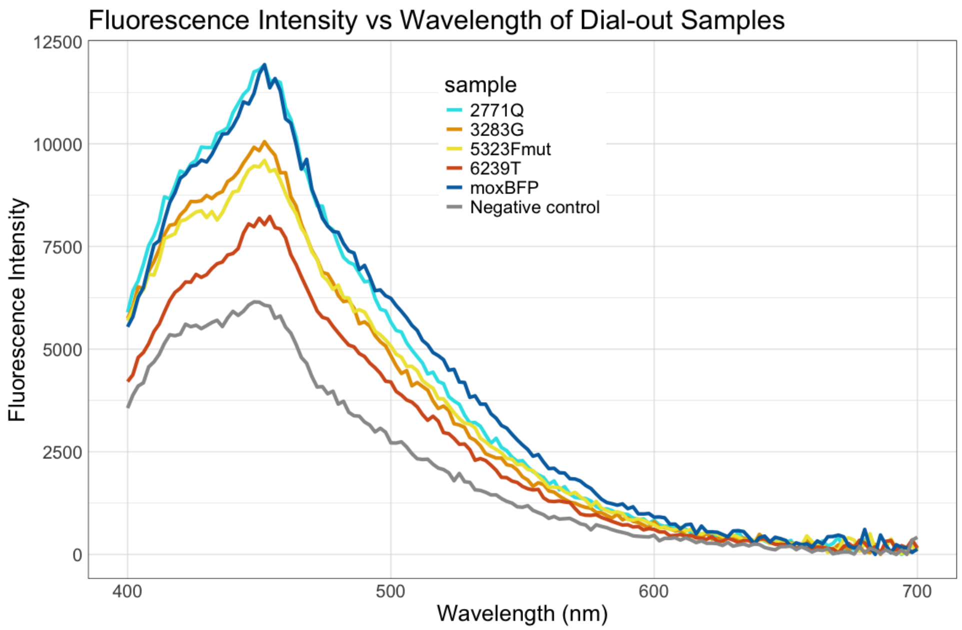

**Fig. S14**. Plate reading of Dial-out samples grown under 18°C culture conditions, using 355 nm excitation and detecting at a fluorescent sweep between 400 and 700. Plots are the average of triplicates normalized to the OD600.

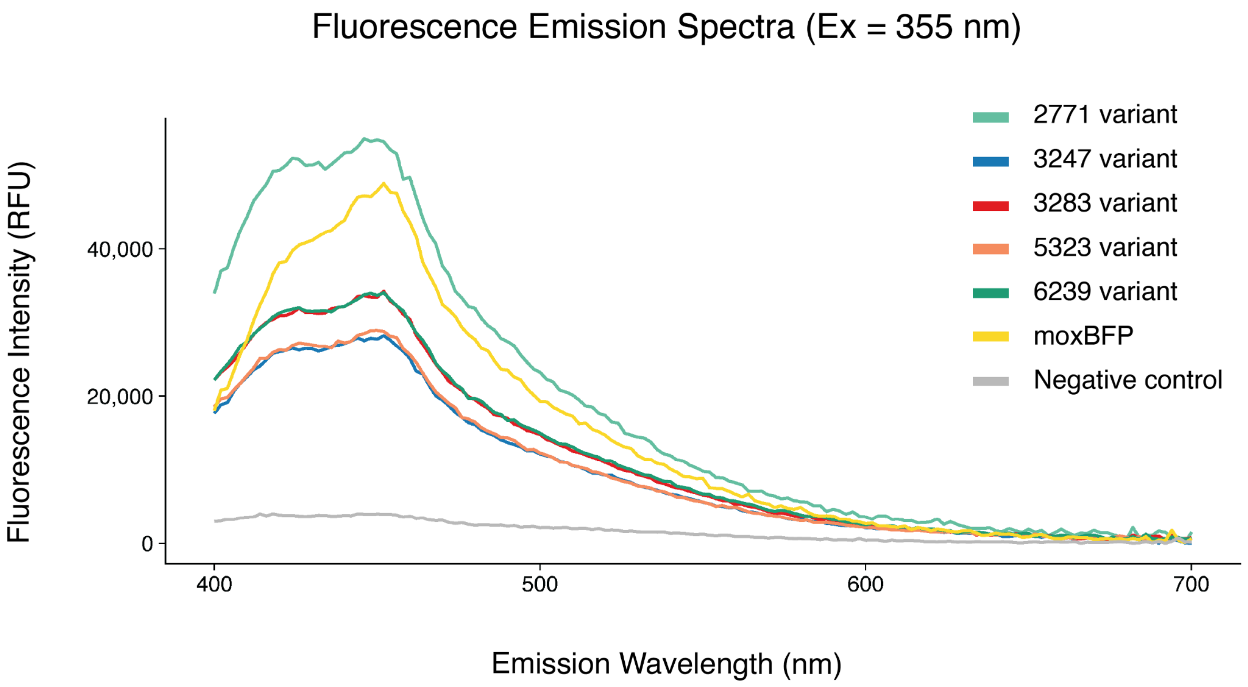

**Fig. S15**. Fluorometer reading of Dial-out samples grown under 18°C culture conditions, using 355 nm excitation and detecting at a fluorescent sweep between 400 and 700. Plots are the average of triplicates normalized to the OD600.

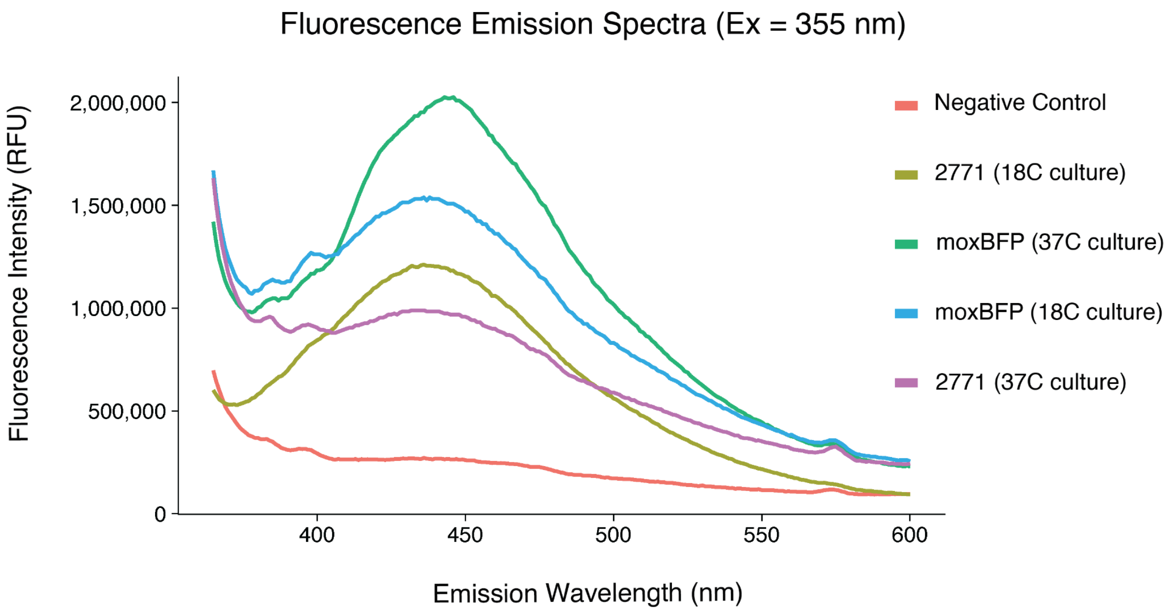

**Fig. S16**. Fluorometer reading of 2771 dial-out sample from two different culture temperatures (18°C and 37°C), using 355 nm excitation and detecting emitted fluorescent intensity between 365 and 600. Plots are the average of triplicates normalized to the OD600 obtained from cuvette reading.

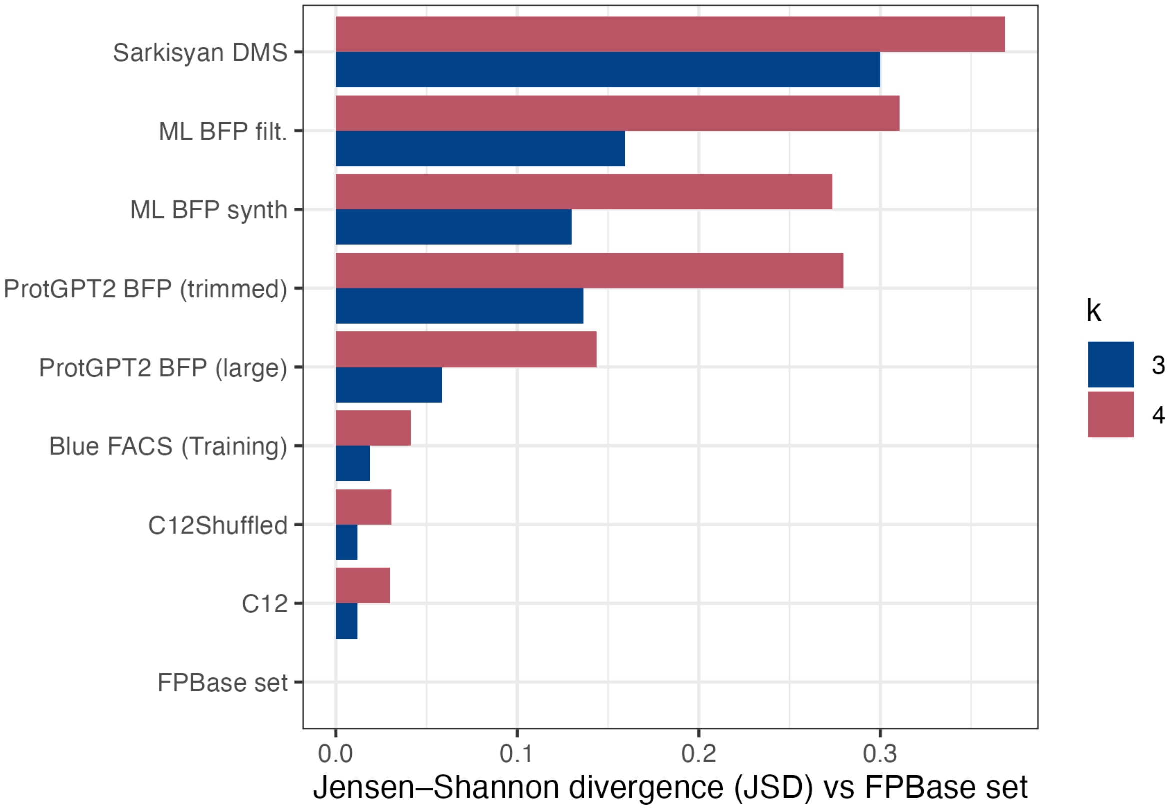

**Fig. S17**. k-mer compositional divergence relative to FPBase. Jensen–Shannon divergence (JSD) between each dataset and the FPBase reference set calculated using amino acid k-mer frequencies for k = 3 (blue) and k = 4 (red). Higher JSD values indicate greater divergence in short-range sequence composition relative to naturally occurring fluorescent proteins. ProtGPT2-generated libraries display substantially higher divergence, particularly at k = 4, indicating remodeling of local sequence motifs.

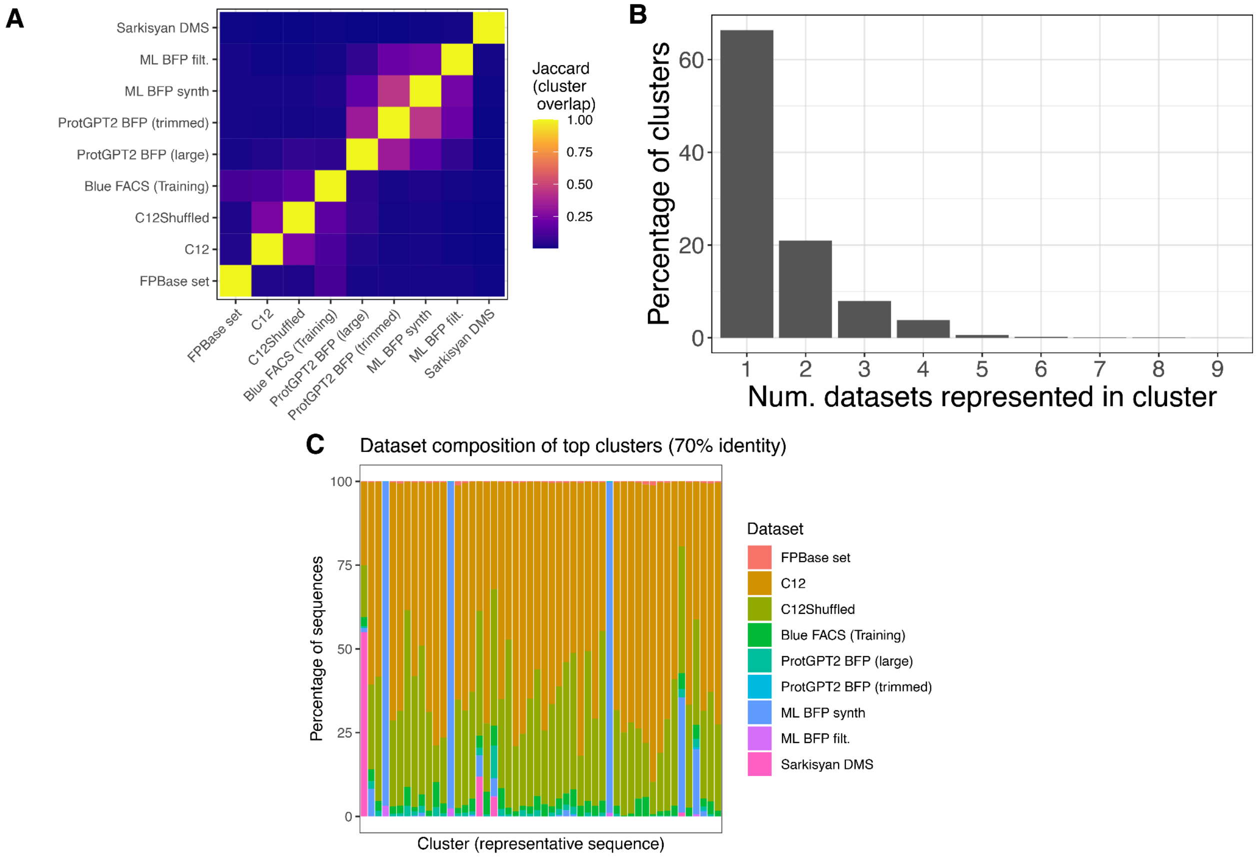

**Fig. S18**. **A.** Cross-set cluster overlap at 70% identity. Deduplicated sequences from all datasets were pooled and clustered at 70% amino acid identity using MMseqs2. Pairwise Jaccard indices were computed based on shared cluster membership. Higher values indicate greater overlap in coarse-grained sequence space occupancy. Parental and shuffled libraries show substantial overlap, while natural FPBase sequences exhibit limited overlap with synthetic and ML-derived datasets. The Sarkisyan GFP DMS dataset remains largely isolated, consistent with its origin as a single-template mutational landscape. **B.** Distribution of dataset representation per cluster. Histogram showing the percentage of clusters represented by one or more datasets at 70% identity. Most clusters are occupied by a single dataset, indicating substantial nonredundant coverage of sequence space. Clusters shared by many datasets are rare, demonstrating that recombination and generative modeling produce distinct regions of occupancy rather than uniformly overlapping ancestral clusters. **C.** Dataset composition of the 50 largest clusters. Stacked bar plot showing dataset contributions to the 50 largest clusters at 70% identity. Each bar represents one cluster, with colors indicating the proportion of sequences contributed by each dataset. Large clusters are frequently dominated by synthetic libraries, with limited contribution from natural FPBase sequences. ML-derived functional sequences are distributed across multiple dominant clusters, indicating retention of structural diversity following experimental validation.

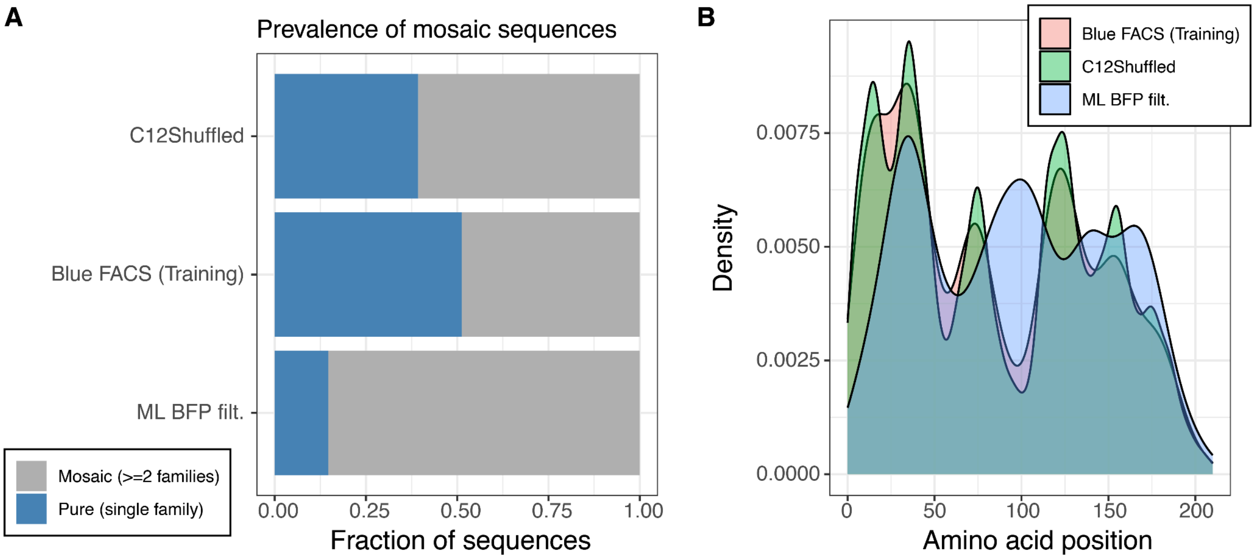

**Fig. S19**. **A.** Prevalence of mosaic sequences across libraries. Stacked bars show the fraction of sequences composed of a single parental family versus those containing segments from two or more families. The ML BFP filt library displays the highest fraction of multi-parental mosaics. **B.** Distribution of recombination boundaries along the fluorescent protein sequence. Density plots show the positions of inferred parental family switches across amino acid coordinates. Shuffled and experimentally selected libraries display localized recombination hotspots, whereas ML-derived variants exhibit broader boundary distributions across the scaffold.

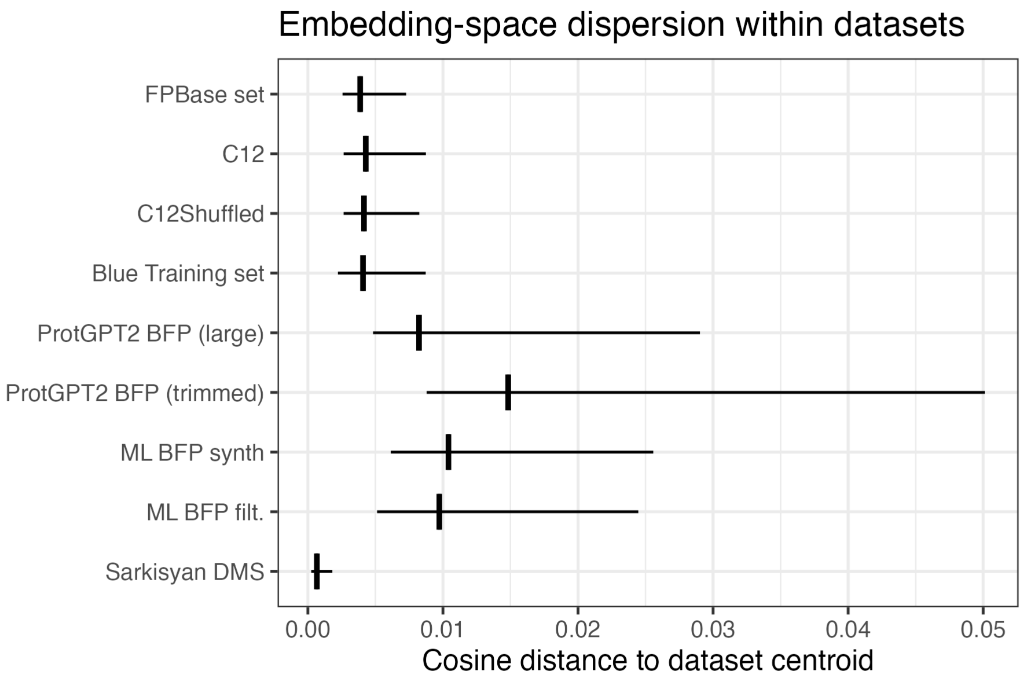

**Fig. S20**. Boxplots of the distributions of cosine distances to each dataset centroid. These show larger within-dataset dispersion for ML-derived libraries compared to FPBase derived sets and DMS variants.

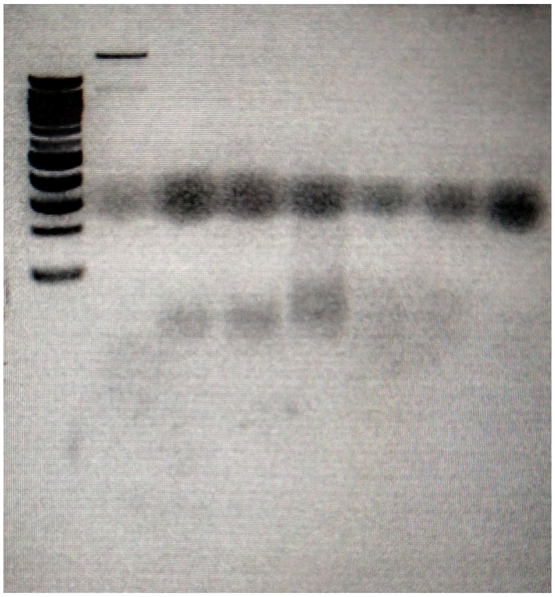

**Fig. S21**. Optimization of DNase I incubation times for 50 uL reactions with 2 ug of linear DNA and 1 uL of 0.25 U/uL DNase I. From left to right the lanes are: 100 bp ladder, no digestion (DNAse added and immediately inactivated at 80C), 1.5min, 1.25min, 1min, 45s, 30s, and 15s of incubation time.

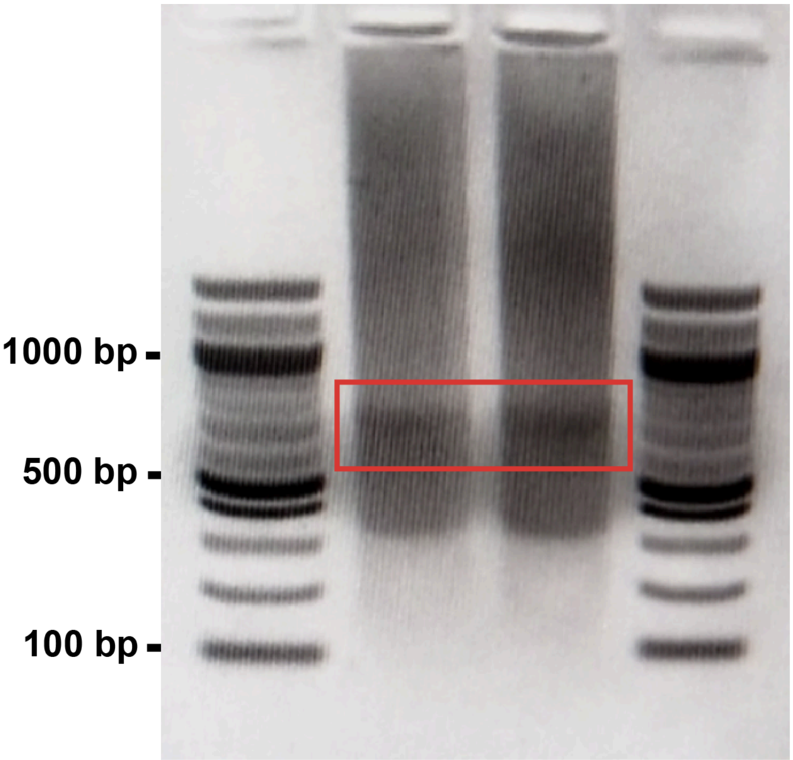

**Fig. S22**. Gel of amplified shuffled library (middle two lanes) with 100 bp NEB ladder for reference (first and last lane). Boxed region is where gel was cut for shuffled library purification.

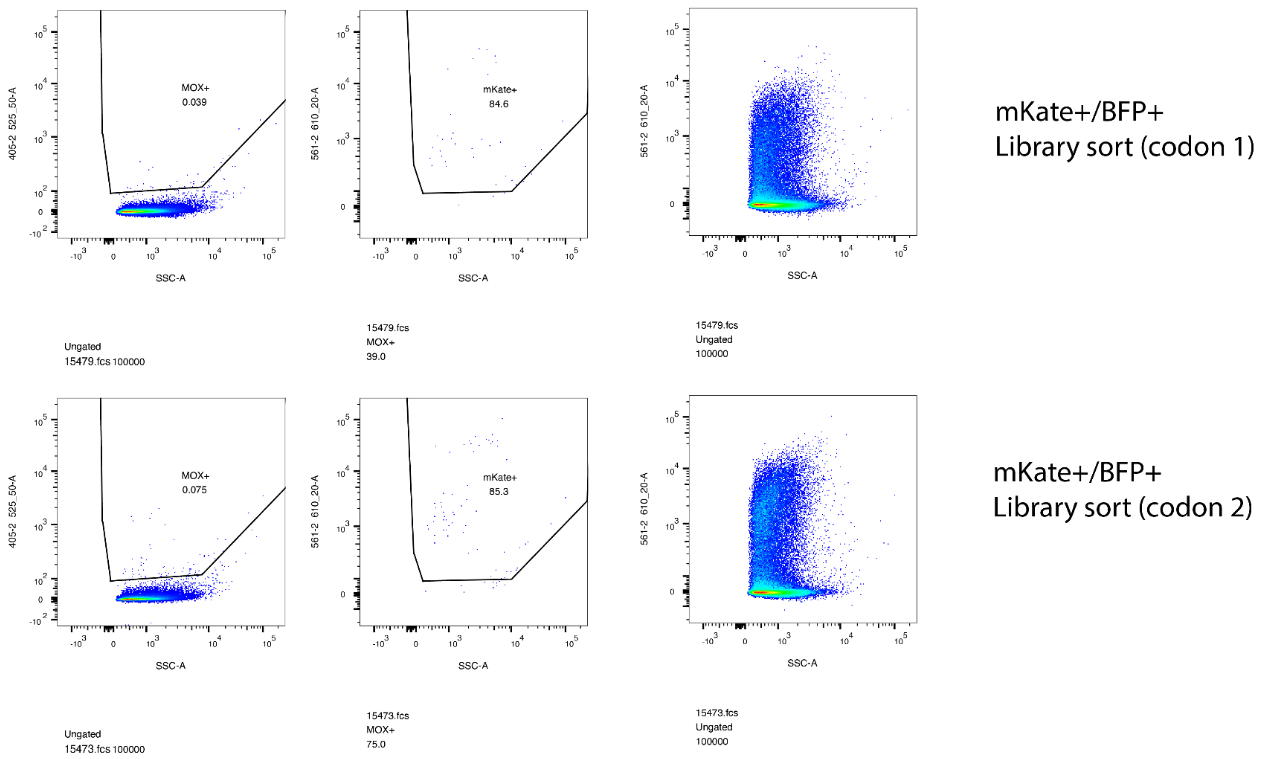

**Fig. S23**. Scatter plots of flow-cytometry gating used to define the mKate⁺/BFP⁺ population in ML-generated E. coli libraries, with representative gating of control samples. Controls included moxBFP (blue), mKate (red), rainbow calibration beads (for fluorescence-to-MFI conversion), and a non-fluorescent E. coli control (no FP plasmid). Cells were gated sequentially on BFP⁺ events (left, 405 laser and 525/50 BP filter), then on mKate⁺ within the BFP⁺ gate (middle, 561 laser, 610/20 BP filter). The right column shows the full, ungated population under red laser/filter set.

**Supplementary Table S1 -** The five dial-out variants isolated and individually characterized in this study.

| **Variant Name** | **Sequence** |
|---|---|
| 6239T | TAKSAVELFTGIVPILVELDGDVHGHKFSVRGEGEGKPYEGTHTVKLQVVEGSPLPFSYDILTTIFQYGNRAFVNYPQDIPDYFKQCFPGGYSWERKFEFEDGGTAIVKSDISLEDGKFIVNVDFKAKDLRRMGPVMQKTIGWDKSFEKMTVSKEVLRGDVTEFLMLEGGGYHSCQFHSTYKPEKPVTLPPNHVVEHQIVRTDLGQSAKGFTVKLEEHAAAHVSL |
| 3283G | GAVNGHKFIIEGDGKGKPFDGTQTMDLTVIEGAPLPFAYDILTTVFDYGNRVFAKYPQDIPDYFKQSFPEGYSWERSMTYEDQGICTATSNISMRGDCFFYDIRFYGTNFPPNGPVMQKKTLKWDPSFEKMTVCDGILKGDVTAFLMLQGGGNYRCQFHTSYKTKKPVTMPPNHVVEHRIARTDLDKGGNSVQLTEHAVAHITSVVPF |
| 3247G | GVSVIFTDKTTVIEGGGKHFSIRSTYEHMRSESDYDSVRRHSTGGSLSGGSRRKLSALTNEDAYAHLVSFINARSSLTDITENVDPHSGEISGVKIRYCELKGVVRLPDYHLVDHCIEILSHDKDYNNVKLYENAVAHSGLPDNARRDNKVQMPKLVEYNSAIHTDLEHHHHHHGSDSDSGSDSDTSRRRHKRRHRHHHHH |
| 2771Q | QTKQPLPISHDIFKKDYTVSSDAKDGSGQSKQSMVTANRFTGGMDELYKSVITSELPVSWTILPWTWCRTTICWPITVRQRPVIQNTTSAVRQLATCCWKITTSCWPITATRSVTTSQATRTCRATTICRRMTPVCWITPALILKTCCCITAATVTSVTARCSTVSRTTPWSMRSVVMSSCHVRRCHRIMTTAAL |
| mut_5323F_f35f5bd7be0d8bcc066a70dedf4989ec6fd944cd748aecf968c9c5524ff9754d | FPLDVTMFLYGNRALTKYPDDIPDYFKQAFPEGLSWERSLEFEDGGSASVSAHISLRGNTFYHKSKFTGVNFPADGPIMQNQSVDWEPNTEKITASDGVLKGDINMALLLEGGGHYRCDFKTTYKAKKVVQLPDYHFVDHHIEIKSHDKDYSNVNLHEHAEAHSELPRQAKDFYVQEKCRCQICTSYQDALVNENIKENMSLKENVVA |

**Supplementary Table S2 -** 4-mer statistics from the k-mer analysis of the datasets.

| Dataset | Total 4-mers | 4-mer richness | Shannon H | N1 |
|---|---|---|---|---|
| FPBase set | 141,647 | 14,865 | 8.215 | 3,696 |
| C12 | 25,059,249 | 91,271 | 8.462 | 4,734 |
| C12Shuffled | 15,085,859 | 68,847 | 8.199 | 3,638 |
| Blue FACS | 4,819,776 | 44,997 | 8.096 | 3,280 |
| Blue Training set | 1,781,291 | 27,455 | 8.071 | 3,202 |
| ProtGPT2 BFP (large) | 1,515,868 | 108,860 | 9.411 | 12,218 |
| ProtGPT2 BFP (trimmed) | 316,655 | 76,411 | 10.185 | 26,500 |
| ML BFP synth | 6,162,338 | 110,993 | 9.218 | 10,076 |
| ML BFP filt. | 137,867 | 28,407 | 8.878 | 7,173 |
| Sarkisyan GFP DMS | 12,638,245 | 15,780 | 5.863 | 352 |

**Supplementary Table S3 -** Primer sequences used in this study.

| Primer name | Sequence | Purpose |
|---|---|---|
| FP Lib 1 Codon 1 FWD | ACCGGTTTCCACGCA | Bulk PCR amplification of Parent library, codon 1 (biotinylated) |
| FP Lib 2 Codon 1 FWD | GGGTTCGAGCGGGAG | Bulk PCR amplification of Parent library, codon 1 (biotinylated) |
| FP Lib 1 Codon 2 FWD | ATCACTCGCGTCCCA | Bulk PCR amplification of Parent library, codon 2 (biotinylated) |
| FP Lib 2 Codon 2 FWD | ACCATCGCGCACCTT | Bulk PCR amplification of Parent library, codon 2 (biotinylated) |
| FP Lib 1 Codon 1 REV | TAGCGCGCAGAGAGG | Bulk PCR amplification of Parent library, codon 1 (biotinylated) |
| FP Lib 2 Codon 1 REV | ACACGCGCGTTGAAG | Bulk PCR amplification of Parent library, codon 1 (biotinylated) |
| FP Lib 1 Codon 2 REV | ACACGCTCGACTCCC | Bulk PCR amplification of Parent library, codon 2 (biotinylated) |
| FP Lib 2 Codon 2 REV | CTCCCTTGCCTGCCC | Bulk PCR amplification of Parent library, codon 2 (biotinylated) |
| pUC19_FWD | ATAAGGGCGACACGGAAATG | |

| | | |
|---|---|---|
| pUC19_REV | CTTTGAGTGAGCTGATACCG | |
| Suppression_501-503-504F | TAAGCGCCCTTCTAATACCCAGGTCTGGCCCTATATACGAATCGGGGATGGTAACTAACG | Suppression primer |
| Suppression_501-503-504R | TAAGCGCCCTTCTAATACCCAGGTCTGGCCCTATATACGAATAGCTGATTGTCCGTTGGT | Suppression primer |
| Ox_01_FWD | AAGAAAGTTGTCGGTGTCTTTGTGTGTGAGCGGATAACAATTTCACACAGGAAACAGCTCATATG | PacBio primer for parent library 1 |
| Ox_01_REV | CGAAAAGTGCCACCTGACGTCGTGCAAGAAAGTTGTCGGTGTCTTTGTG | PacBio primer for parent library 1 |
| Ox_02_FWD | TCGATTCCGTTTGTAGTCGTCTGTTGTGAGCGGATAACAATTTCACACAGGAAACAGCTCATATG | PacBio primer for parent library 2 |
| Ox_02_REV | ACAGACGACTACAAACGGAATCGAGCACGACGTCAGGTGGCACTTTTCG | PacBio primer for parent library 2 |
| Mkatelinker_plasmid_rev | TCGCTCACCATagctgtttcctgtgtgaaattgttatcc | has overhang that is complementary to Mkate_nondeI_for |
| Mkatelinker_plasmid_for | ACCttgaaatgaacctaaGTGTGGCTGCGGAACcat | forward, has overhang that is complementary to linker fragment |
| Mkate_nondel_for | agctATGGTGAGCGAACTGATTAAAGAAAACAT | agctATGGTGAGCGAACTGATTAAAGAAAACAT |
| mkate_nondel_rev | AGGTTTCTTTATCCGCTTCTTTAATACGTTCA | Complementary to 3' end of mkate and linker fragment |
| skpp504R | ATAGCTGATTGTCCGTTGGT | Used for Q5 PCR-amplifying chimeric genes after DNA shuffling |
| skpp504F | ATCGGGGATGGTAACTAACG | Used for Q5 PCR-amplifying chimeric genes after DNA shuffling |
| pUC19 FWD | ATAAGGGCGACACGGAAATG | Used to validate shuffled genes via colony PCR |
| pUC19 REV | CTTTGAGTGAGCTGATACCG | Used to validate shuffled genes via colony PCR |
| EVBC8_FWD1 | aggAGAAGAGCGCACGACGTCACGTCGCAGAATTCCTTTTCGGGGAAATGTGCGCGGAAC | Around the horn PCR primer for ml library backbone |
| mKateBC_REV1 | gccgtCATATGTcCTCGAGCTGCAGCTTT | Around the horn PCR |

| | | |
|---|---|---|
| | GGCA | primer for ml library backbone |
| EVBC8_FWD2 | gtgGGTACCtaaGTGTcGCTGCcGAACagcNNNNNNNNNNNNNNNNNNNNaggAGAAGAGCgcacGACGTcaCg | PCR2 - adding barcodes to ml library backbone |
| mKateBC_REV2 | biotin -gccgtCATATGTcCTCGAGCTGCAGCTT | PCR 2 - adding barcodes to ml library backbone; also used in PCR 3 |
| EVBC3_FWD3Re | biotin -GTGGGTACCTAAGTGTCGCTGCCGAACAG | Amplifying barcoded backbone |
| skpp15-1-F filt15-1651 | GGGTCACGCGTAGGA | Amplification of oligo subpools using qPCR (biotinylated; ML-generated library) |
| skpp15-1-R filt15-1181 | GTTCCGCAGCCACAC | Amplification of oligo subpools using qPCR (biotinylated; ML-generated library) |
| skpp15-2-R filt15-1539 | GCCGTGTGAAGCTGG | Amplification of oligo subpools using qPCR (biotinylated; ML-generated library) |
| skpp15-2-F filt15-568 | CGCGTCGAGTAGGGT | Amplification of oligo subpools using qPCR (biotinylated; ML-generated library) |
| 6239T REV | GATATCATATGACCGCCAAGAGTG | Reverse dial out primer for variant 6239T |
| 6239T FWD | ACAATGGTTGCGTGCGCAAC | Forward dial out primer for variant 6239T |
| 584auto REV | GATATCATATGGCTCCGAGC | Reverse dial out primer for variant 584auto |
| 584auto FWD | CTCCTTAAAGTGTTGACTGTCCTAT | Forward dial out primer for variant 584auto |
| 3618H REV | GATATCATATGcattggccgactacccacgg | Reverse dial out primer for variant 3618H |
| 3618H FWD | TCTCCTTATGTTCATATAGGAATGAG | Forward dial out primer for variant 3618H |
| 3283G REV | GATATCATATGGGTGCTGTAAATGGC | Reverse dial out primer for variant 3283G |
| 3283G FWD | TCTCCTGAAATGTACCAAGCATTTAC | Forward dial out primer for variant 3283G |
| mut_7109V REV | GATATCATATGGTTATGGGTGACGT | Reverse dial out primer |

| | | |
|---|---|---|
| | | for variant 7109V |
| mut_7109V FWD | GTCCATGGCTCCATAACCAT | Forward dial out primer for variant 7109V |
| mut_5323F REV | GATATCATATGTTTCCGCTGGACGTA | Reverse dial out primer for variant mut_5323F |
| mut_5323F FWD | CTGTGGCCCTATTAGAGTAAGG | Forward dial out primer for variant mut_5323F |
| 3247G REV | GATATCATATGGGCGTTAGTGTAATCTTC | Reverse dial out primer for variant 3247G |
| 3247G FWD | TCTCCTAACATAGCTTAGATATAAGG | Forward dial out primer for variant 3247G |
| 7255V REV | GATATCATATGgtatacgagggcgtacaaact tggccag | Reverse dial out primer for variant mut_7255V |
| 7255V FWD | TCTTCTCCTAAAAACTAAATCACAATTGA | Forward dial out primer for variant mut_7255V |
| mut_6473W | GATATCATATGTGGATGGTAGATCTGAC | Reverse dial out primer for variant mut_6473W |
| mut_6473W FWD | TCTCCTTGTCTTTTATCAAAAGAAAT | |
| 2771Q REV | GATATCATATGCAAACGAAGCAACCT | Reverse dial out primer for variant 2771Q |
| 2771Q FWD | CCTCTTAACGGACAAGCGAAAAA | |
| 4379K REV | GATATCATATGAAGGCTTGGGG | Reverse dial out primer for variant 4379K |
| 4379K FWD | TCTCCTAGCAATGAGTAAAGATAATC | |
| Suppression_501-503-504F | TAAGCGCCCTTCTAATACCCAGGTCTGGCCCTATATACGAATCGGGGATGGTAACTAACG | Used in ePCA step for DropSynth |
| suppression_501-503-504R | TAAGCGCCCTTCTAATACCCAGGTCTGGCCCTATATACGAATAGCTGATTGTCCGTTGGT | Used in ePCA step for DropSynth |
| skpp503F | AGGTCTGGCCCTATATACGA | Used for Suppression PCR at the end of th DropSynth protocol |
| Linker fragment | CGGATAAAGAAACCTACGTTGAACAACATGAAGTCGCTGTGGCCCGTTACTGCGATCTGCCGTCCAAGCTCGGCCATCGAGGCAGCCTGGCGGAAGCAGCGGCCAAAGAGGCGGCAGCTAAGGAAGCGGCTGCCAAAGCTGCAGCTCGAGGaGTGcatATg*gctttc*GGTACCttgaaatgaacctaa | Fragment that includes a portion of mKate, the alpha helix linker, and ndeI/kpnI restriction sites: |
| pEVBC_FWD | gccgtCATATGagctgtttcctgtgtgaaattg | Used to amplify pEVBC1 PCR2 |
| pEVBC_amp_FWD | gtgGGTACCtaaGTGTGGCTGCGGAAC | Used to amplify pEVBC1 PCR2 |
